# The Alzheimer’s disease risk gene *SORL1* is a regulator of excitatory neuronal function

**DOI:** 10.1101/2025.07.28.667194

**Authors:** C. Andrew Williams, Shannon E. Rose, Vera Stamenkovic, Stephen E.P. Smith, Jessica E. Young

## Abstract

**Background:** Synaptic dysfunction is an early feature of Alzheimer’s disease (AD) and a significant contributor to cognitive decline and neurodegeneration. Proper localization of proteins involved in pre-and post-synaptic composition is dependent on endosomal recycling and trafficking. Alterations in trafficking complexes, such as retromer, have been shown to impair neuronal synaptic function. The *SORL1* gene has been strongly implicated in AD pathogenesis and its protein product, SORLA, is an endosomal receptor that works in conjunction with retromer to regulate endosomal recycling.

**Methods:** We utilized our established human induced pluripotent stem cell (hiPSC) derived excitatory cortical neuron model to examine *SORL1*’s role in synaptic protein composition and neuronal function. We used Quantitative Multiplex co-Immunoprecipitation (QMI), a mesoscale proteomics assay to measure synaptic protein interactions, immunocytochemistry to assay synapses and AMPA receptor subunits, and multi-electrode arrays (MEAs) to measure neuronal function of *SORL1* KO and isogenic control hiPSC derived neurons.

**Results:** We show that loss of *SORL1* expression significantly changes many synaptic protein-protein interactions and patterns of expression. We demonstrate that *SORL1* deficient neurons are hyperactive and that the increased activity is driven by glutamatergic neurotransmission. Hyperexcitability has been seen in other models of AD with familial AD variants in amyloid precursor protein and presenilin genes, due to the increases in amyloid beta (Aβ) peptides. In the case of *SORL1* deficiency, the hyperexcitability we observe is primarily due to mis-trafficking of synaptic proteins, rather than an overall increase in Aβ. Finally, we find that *SORL1* deficient neurons have impaired synaptic plasticity.

**Conclusions:** These findings further support a growing body of literature implicating early endosomal recycling defects as drivers of AD pathogenesis. Furthermore, our work supports further emphasis on exploring the SORL1-retromer pathway for therapeutic development in AD.

## Background

Alzheimer’s disease (AD) is characterized by progressive synapse loss and dysfunction that precedes widespread neuronal death and correlates strongly with cognitive decline (Tzioras 2023). Synaptic alterations include changes in synapse morphology, impaired synaptic plasticity and disrupted neurotransmission, particularly affecting glutamatergic neurons in the cerebral cortex (Griffiths and Grant 2023). Hallmark AD pathogenic proteins amyloid beta (Aβ) and hyper-phosphorylated tau (pTau) interfere with synaptic function by impairing long-term potentiation and disrupting synaptic transmission (John and Reddy 2021) but how AD risk genes impact synapses is largely unknown.

Pathogenic changes to neuronal synapses and disruption of signaling can lead to aberrant neuronal activity and larger scale network activity dysfunction. Neuronal hyperactivity is associated with early disease state, followed by hypoactivity with advancing disease status. Disease-associated hyperactivity can manifest as an increased prevalence of seizures and epileptiform brain activity (Vossel 2017) and is associated with faster rates of cognitive decline. Neuronal hyperexcitability is seen in multiple models of AD (Targa Dias Anastacio 2022) and has emerged as a significant contributor to synaptic dysfunction. This hyperexcitability manifests as increased spontaneous neural activity, altered calcium homeostasis and enhanced susceptibility to excitotoxicity (Targa Dias Anastacio 2022), altered glutamate receptors, and changes to excitatory-inhibitory balance (Lerdkrai 2018, Ghatak 2019). Ultimately, this cascade of events can lead to synaptic exhaustion and degeneration through excessive metabolic demands and increased cellular stress (Katsu-Jimenez 2017, Guo 2017).

Deficiencies in endosomal recycling can contribute to synaptic dysfunction. Reduced expression of the retromer complex, a multiprotein complex heavily involved in endosomal trafficking and recycling, lead to loss of cell-surface glutamatergic AMPA receptors and impaired long-term potentiation (Temkin 2017, Simoes 2021). The sortilin-related receptor SORLA (*SORL1)* directly interacts with retromer to facilitate endosomal through the recycling, retrograde, and degradative pathways (Barthelson 2020). *SORL1* is an established AD risk gene, implicated by multiple GWAS studies (Kim 2025, Bellenguez 2017, Lambert 2013) and by exome studies showing that loss of function variants in SORL1 are causal for early onset AD (Holstege 2017). Furthermore, SORLA protein levels are decreased in the AD human brains and specifically decreased in the AD-susceptible *trans-*entorhinal cortex (Simoes 2021). We have previously established models of *SORL1* deficiency using human induced pluripotent stem cell (hiPSC)-derived neurons and glia (Knupp 2020, Mishra 2022, Mishra 2025). In cortical neurons with loss of *SORL1* expression we have observed altered trafficking of key proteins involved in neuronal synaptic function including the GLUR1 subunit of the AMPA receptor and the BDNF receptor TRKB (Mishra 2022). However, how *SORL1* expression shapes neuronal function and synapses remains underexplored.

Here, we used *SORL1* deficient human cortical neurons to investigate the role *SORL1* plays in synaptic structure and neuronal activity. We report that *SORL1* KO neurons have disrupted synaptic protein interactions and alterations of excitatory postsynaptic proteins and synapses. We find that excitatory glutamate AMPA receptor subunits have altered expression profiles in *SORL1* KO neurons. We observe that *SORL1* deficient neurons are hyperexcitable relative to isogenic wild-type neurons and that the increased neuronal activity is driven principally by glutamate excitatory neurotransmission. This hyperexcitability phenotype is not solely due to increased secretion of soluble Aβ peptides. We also demonstrate the *SORL1* deficient neurons have impaired long-term potentiation, demonstrating, for the first time, that *SORL1* plays a critical role in activity-driven modification of the synapse. Together, our study shows that our human neuron model of *SORL1* deficiency recapitulate an early phenotype of AD pathology that may represent a novel therapeutic target.

## Methods

### Cell lines and CRISPR-Cas9 genome editing

All cell lines used in this study have been previously published (Knupp 2020, Mishra 2022, Mishra 2023, Shin Evitts 2023). All cell lines used are isogenic and generated from the published and characterized CV cell line (Young 2015, Levy 2007). This cell line is male with an APOE e3/e4 genotype. CRISPR-Cas9 gene editing was used to generate *SORL1* KO clones and APP^Swe+/+^ as described previously (Knupp 2020, Young 2018). All clones demonstrated normal karyotypes and regularly tested for mycoplasma (MycoScope, AMS Bio).

### Neuronal differentiation

hiPSCs were differentiated into neurons using dual-SMAD inhibition (Rose 2018, Knupp 2020, Mishra 2022, Mishra 2023). Briefly, hiPSCs were plated on Matrigel (Corning cat. #356231) coated culture plates and expanded with mTesR Plus (StemCell cat. #100-0276) until near-confluent. On day 1 of neural induction, cell medium was switched to basal neural maintenance medium (BNMM) with dual-SMAD inhibitors, composed of: 1:1 DMEM/F12 + glutamine (Gibco cat. #11320-033) and Neurobasal (Gibco cat. #21103-049), supplemented with 1% B27 (Gibco cat. #17504-044), 0.5% N2 (Gibco cat. #17502-048), 0.5% GlutaMax (Gibco cat. #35050-061), 0.5% insulin-transferrin-selenium-A (Gibco cat. #51300-044), 0.5% Non-essential amino acids (NEAA) (Thermo Fisher cat. #11140050), 0.2% β-mercaptoethanol (Thermo Fisher cat. #21985023), and dual-SMAD inhibitors 10μM SB-431542 (Biogems cat. #BG6675SKU301) and 0.5μM LDN-193189 (Biogems cat. #BG5537SKU106). Cells were fed daily for seven days. On day eight, cells were dissociated and split with Versene (Gibco cat. #15640066). One day after passaging, medium was switched to BNMM supplemented with 20ng/ml fibroblast growth factor (FGF) (R&D Systems cat. #233-FB/CF). Cells were passaged and split on day sixteen, continuing to be fed with BNMM+FGF. On approximately day twenty-three, cells were sorted using fluorescence-activated cell sorting (FACS) flow cytometry. Quad-stained cells were isolated for CD24/CD184 positivity and CD44/CD271 negativity(Yuan 2011) to isolate neural precursor cells (NPCs). After FACS, NPCs were expanded for neural differentiation. Eight million NPCs were plated on Matrigel-coated 10cm culture plates. After twenty-four hours, medium was switched from BNMM+FGF to BNMM + 0.2 μg/ml brain-derived neurotrophic factor (BDNF) (PeproTech cat. #450-10), 0.2 μg/ml glial-derived neurotrophic factor (GDNF) (Peprotech cat. #450-02), and 500 μM db-cAMP (Sigma-Aldrich #D0627). Media was changed twice a week for three weeks. After three weeks of neuronal differentiation, cells were dissociated with Accutase (Innovative Cell Technologies cat. #AT-104). Cell suspension concentrations were counted using Trypan blue (Gibco cat. #15250-061) and a TC20 cell counter (Bio-Rad). Cells were then replated according to experimental needs. All cells were maintained at 37° C and 5% CO_2_.

### Quantitative Multiplex Immunoprecipitation (QMI)

Following differentiation, neurons were passaged to multiple-electrode arrays (see below) and seven million cells were plated on parallel Matrigel-coated 6 well culture plates. In the first week post plating, cultures were transitioned from BNMM medium containing BDNF, GDNF, and db-cAMP (see above) to Brainphys neuronal medium (Stemcell Technologies cat. #05790) supplemented with 1% B27, 0.5% N2, 0.2 μg/ml BDNF, 0.2 μg/ml GDNF, 500 μM db-cAMP, and 1 μg/ml laminin (Invitrogen cat. #23017-015). Two-thirds of well volume was changed twice a week. MEA plates were monitored for the development of network bursting activity. Upon network bursting detection, approximately 30 days post plating (DPP) after neuronal differentiation, parallel 6 well cultures were harvested for QMI analysis.

Multiplexed immunoprecipitation is achieved using antibody-based immunoprecipitations onto color-coded Luminex beads, followed by probing for co-associated proteins using fluorophore-coupled antibodies and quantification on a flow cytometer, as described previously (Lautz, 2018, 2021; Heavner 2021). Briefly, samples were lysed in NP40 lysis buffer [150mM NaCl, 50mM Tris (pH 7.4), 1% NP-40, 10 mM NaF, 2 mM sodium orthovanadate + Protease/phosphatase inhibitor cocktails (Sigma)] and protein concentrations were measured by BCA assay (Pierce) and normalized between samples. A master mix containing 20 classes of antibody-coupled Luminex beads was prepared and added to lysates. Samples were incubated overnight at 4 °C on a rotator. The following day, beads were washed with cold FlyP buffer [50 mM tris (pH7.4), 100 mM NaCl, 1% bovine serum albumin, and 0.02% sodium azide] and distributed into 40 wells of a 96-well plate for technical replicates. One of 20 biotinylated probe antibodies was added to each well, in duplicate, and the plate was incubated at 4 °C with gentle agitation for 1 hour. After incubation, the resulting bead-probe complexes were washed three times using an automatic plate washer with cold FlyP buffer, incubated with streptavidin-phycoerythrin at 4 °C for 30 min with gentle agitation, and washed an additional three times. Finally, samples were resuspended in 125 μl of ice-cold Fly-P buffer and bead fluorescence was measured from a mean of 60 beads/class/well using a customized refrigerated Bio-Plex 200. For comprehensive video protocol, analysis software code for both R and MATLAB, refer to Brown et al 2019. QMI validation, including specific validation of each antibody pair, was described previously (Lautz 2018). The full list of IP_ probe antibody pairs used in the study is provided in Table S1.

Raw data from each well were processed using the BioPlex200’s built-in software to (i) eliminate bead doublets based on doublet discriminator intensity thresholds (>5000 and <25,000 arbitrary units; Bio-Plex 200), (ii) identify specific bead classes according to defined bead regions, and (iii) match individual bead phycoerythrin fluorescence measurements to their corresponding bead regions. This processing generated a distribution of fluorescence intensity values for each pairwise protein interaction measurement. XML output files were parsed to acquire the raw data for use in MATLAB using the ANC program (Smith 2016).

### ANC/CNA analysis

Data from two experiments were input into the ANC program (Smith 2016) as individual batches (N of 4). ANC uses a modified non-parametric Kolmologorv-Smirnoff test to identify interactions that are significantly different between control and *SORL1* KO neurons in >70% experiments at a Bonferroni-corrected p<0.05. Median fluorescent intensity matrices were then exported to R, ComBAT-normalzied (SVA package CIte) and input into a custom CNA script in R (Horvath and Langfelder 2008, Brown 2019 (as above). Artificial variance (<1%) was added for 27 interactions with 0 variance to allow ComBAT to include them, which resulted in no experiment-level batch effects. CNA modules were identified using a soft power of 8 and minimum module size of 10 with a module merge cut-off of 0.4. Modules with a p > 0.05 correlation for the “genotype” variable were considered significant. Interactions that met two independent statistical criteria of 1) individually ANC significant and 2) module membership (p<0.05) in a CNA module that was significantly (p<0.05) associated with the experimental variable “genotype”, were reported in Fig 1. The heatmap in Fig 1C was generated with the heatmap.2 command in R, and bar graphs in Fig 1D-I were generated using combat-normalized MFIs in Prism (Graphpad).

**Figure 1:**
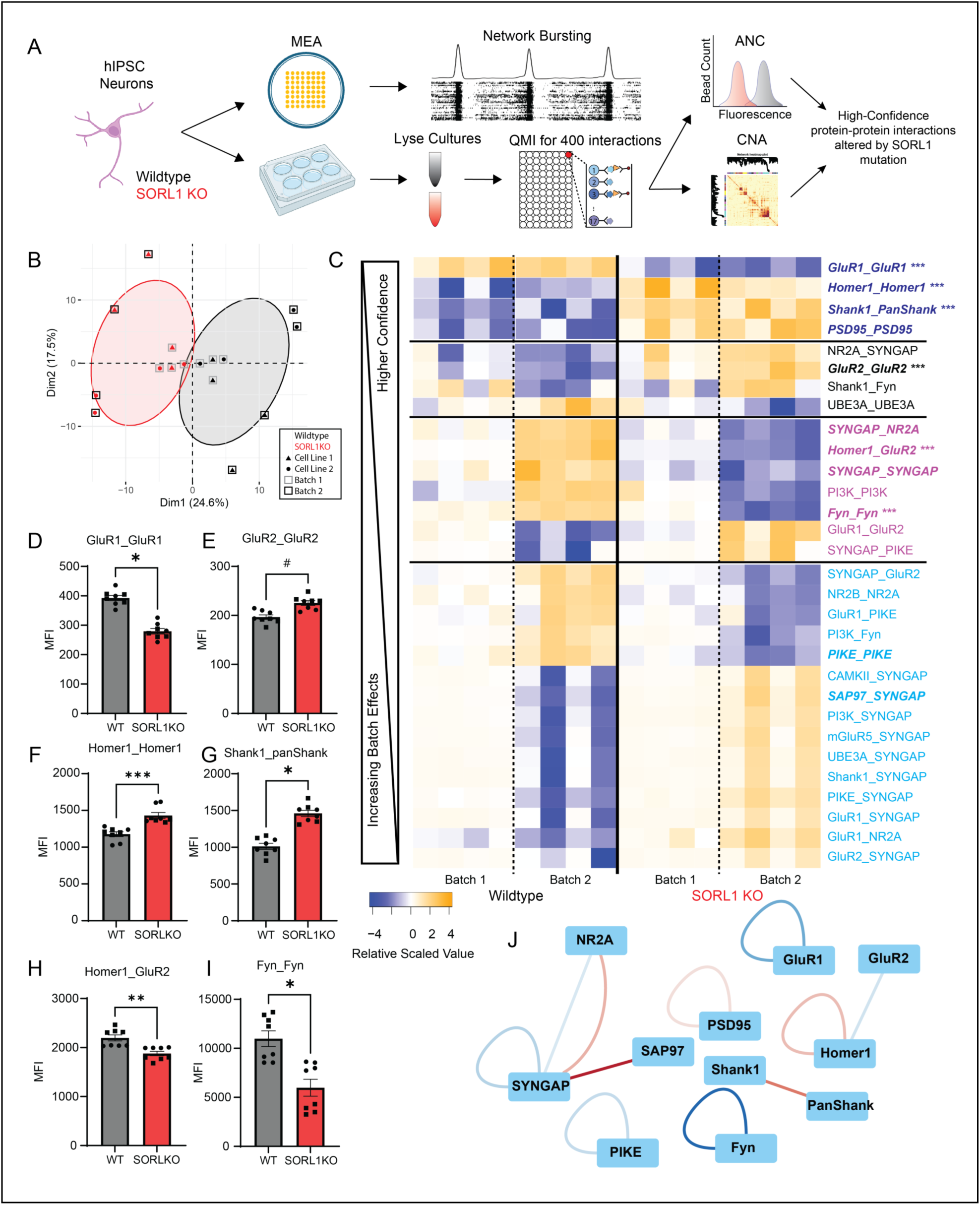
Widespread alterations in the *SORL1* KO glutamate synapse interactome. A) Experimental design. B) Principal component graph of QMI data matrices. C) Heatmap of interactions significantly altered in human *SORL1* KO neurons. Each row represents an interaction that was significant by ANC in at least one batch, each column represents a biological replicate. Interaction color indicates the module with which the interaction is most strongly correlated. Bold-italic interactions were significant across both batches. *** indicate interactions shown in D-I. D-I) Median fluorescent intensity (MFI) is shown for 6 example interactions. Circles indicate batch 1, squares batch 2. * indicates ANC-significant (p<0.05) for both batches, # for only batch 2. J) Node-edge diagram showing interactions that were significantly increased (red) or decreased in both batches of human *SORL1* KO neurons.

### Immunocytochemistry

Following differentiation, neurons were passaged to Matrigel-coated #1 glass coverslips in 24 well culture plates. Neurons were plated at a density of 500,000 cells per coverslip. In the first week post plating, cultures were transitioned from BNMM medium containing BDNF, GDNF, and db-cAMP (see above) to Brainphys neuronal medium supplemented with 1% B27, 0.5% N2, 0.2 μg/ml BDNF, 0.2 μg/ml GDNF, 500 μM db-cAMP, and 1 μg/ml laminin. Two-thirds of well volume was changed twice a week. At approximately DPP 30, cell medium was removed, coverslips were washed with Dulbecco’s modified phosphate-buffered saline with calcium and magnesium (DPBS+) (Cytiva cat. #SH30264.01) and immediately fixed in 4% paraformaldehyde (PFA) for 15 minutes at room temperature. Following fixation, coverslips were washed three times with DPBS+ to remove PFA. Cells were blocked and permeabilized with 5% normal goat serum (Jackson Laboratories cat. #005-000-121), 2.5% bovine serum albumin (Sigma-Aldrich cat. #A7906), and 0.1% Triton X-100 (Fisher cat. #BP151) for one hour at room temperature. Cells were immediately incubated with primary antibodies at 4°C overnight. HOMER1 (Synaptic Systems cat. #160003 1:250), SYNAPSIN1/2 (Synaptic Systems cat. #106004 1:500), GLUR1 (Millipore cat. # MAB2263 1:500), GLUR2 (ProteinTech cat. #11994-1-AP 1:500), MAP2 (Abcam cat. #ab92434 1:1000). Following primary antibody incubation, coverslips were washed three times with PBS+ with 0.1% Triton X-100. Cells were then stained with secondary antibodies for one hour at room temperature. Secondary antibodies were conjugated to AlexaFluor fluorophores 488, 594, or 647 (Invitrogen) and used at a dilution of 1:1000. Cells were then washed three times with PBS+ to remove unbound secondary antibodies and coverslipped with ProLong Gold mounting medium containing DAPI stain (Invitrogen cat. #P36931) and sealed prior to confocal imaging.

### Confocal microscopy and image analysis

Cells were imaged using a Leica SP8 confocal microscope with a 63x oil immersion objective. All images were captured with the experimenter blinded to cell line. All images were captured and subsequently deconvolved using the Leica LIGHTNING deconvolution method using the Adaptive mode. Captured images had a resolution of 1040 × 1040 pixels with 2x zoom. Laser lines were utilized sequentially. Laser power and detector settings are as follows. For HOMER1 and SYNAPSIN synaptic staining, 405nm (1.00% 80% HyD gain), 488nm (2.00% 800V PMT gain), 552nm (2.00% HyD 243% HyD gain), 638nm (0.1% 840V PMT gain). For GLUR1 and GLUR2 characterization, 405nm (1.00% power, 185% HyD gain), 488nm (1.5% power, 773V PMT gain), 552nm (2.00%, 450% HyD gain), 638nm (0.7% power, 576V PMT gain) settings were used. Z-stacks of synapse puncta were captured with a total Z-size of 5.5 µm in 12 total steps of 0.5 µm. Z-stacks of GLUR1/GLUR2 stained neurons were captured with a total Z-size of 3.25 µm in 17 total steps of 0.2 µm. All imaged neurons were located by MAP2 and DAPI signals.

Image analysis of HOMER1/SYNAPSIN and GLUR1/GLUR2 immunostained neurons was performed using Bitplane Imaris software (Oxford Instruments). Similar pipelines were created for both synaptic and glutamate receptor analyses, as follows: The surface object workflow was used to define the MAP2+ neuron regions of interest (whole cell, soma, dendrites) and for the creation of masked HOMER1/SYNAPSIN and GLUR1/GLUR2 channels within the neuron ROIs. The surface object workflows were then created to quantify HOMER1/SYNAPSIN and GLUR1/GLUR2 puncta volume, number, and fluorescent intensity. Colocalization of HOMER1 and SYNAPSIN was analyzed using the ImarisColoc function.

### Multiple electrode array (MEA)

Differentiated neurons were generated as described above and were passaged to Matrigel-coated MEA culture plates in high-density droplets. 6-well and 48-well MEA plates were used for these experiments (Axion Biosystems cats. #M384-tMEA-6W and #M768-tMEA-48W). Matrigel coatings were limited to the area of the recording electrodes. Upon plating, Matrigel droplets were aspirated and neurons were plated on the recording electrodes at a density of 10,000 cells/µl. 300,000 total cells were plated on 6-well MEA plate wells and 80,000 total cells on 48-well MEA plate wells. In the first week post plating, cultures were transitioned from BNMM medium containing BDNF, GDNF, and db-cAMP (see above) to Brainphys neuronal medium supplemented with 1% B27, 0.5% N2, 0.2 μg/ml BDNF, 0.2 μg/ml GDNF, 500 μM db-cAMP, and 1 μg/ml laminin. Two-thirds of well volume was changed twice a week.

### MEA recordings and analysis

Extracellular electrical activity of hiPSC neurons plated on MEA plates was recorded twice a week using Axion Biosystems Maestro Pro system. Recordings were taken prior to medium changes to minimize physical perturbation of neurons. MEA culture plates were acclimated to the MEA reader for approximately ten minutes prior to the onset of recording and the environment was controlled at 37° C and 5% CO_2_. Recordings were approximately four minutes in duration. Signals were recorded at a sampling frequency of 12.5 kHz with a 3 kHz Kaiser window low-pass filter and a 200 Hz high-pass filter. Spikes were detected using the adaptive threshold crossing method of the Axion Axis Navigator software. Electrodes were not included in downstream analyses if they failed to detect more than five spikes a minute. All MEA metrics were calculated from a three-minute length of recording. Mean firing rate and coefficient of variation of the inter-spike interval metrics were calculated using the Axion Neural Metric Tool software. Network bursts were detected and metrics quantified using custom Matlab (Mathworks) scripts in the following manner. Spike times per electrode were first imported into Matlab from Axion .spk files. Spike times across all electrodes were combined and binned with bin sizes of 200ms. Spike time histogram plots of the binned spike times were calculated for each well and a gaussian filter was applied. Network bursts were identified as peaks of the spike time histogram that surpassed an adaptive threshold that was determined by the standard deviation of the overall levels of activity on spike histogram and a minimum spike histogram peak value of thirty action potentials. The start and end timepoints of network bursts were detected by the first intersections of the spike histogram before and after a detected peak and a threshold determined by the median spiking value of the recording. Detected network bursts that did not have spiking activity detected on 20% of electrodes were then discarded. Network bursting, intraburst spiking, and extraburst spiking metrics were then quantified using the start, peak, and end timepoints identified from the spike histogram.

### Neurotransmitter receptor antagonist experiments

Blockade of AMPA, NMDA, and GABA_A_ neurotransmitter receptors were performed pharmacologically with the addition of NBQX (Tocris, cat. #0373), D-AP5 (APV, Tocris cat. #0106), and SR 95531 hydrobromide (Gabazine, Tocris cat. #1262), respectively. A full media change was performed the day before antagonist application. Antagonists were diluted in Brainphys medium at 10x the final concentration. Baseline recordings were performed as described above. After determination of baseline activity, 10x antagonist-containing medium was added to each MEA well for a final antagonist concentration of NBQX 10 µM, APV 50 µM, or Gabazine 10 µM. Recordings were then immediately taken and wells were washed with three full volume medium changes of Brainphys neuronal medium. Approximately two hours later, washout condition recordings were performed.

### Chemical long-term potentiation (cLTP)

cLTP methods were adapted from Pré et al 2022. Following maturation and development of network bursting, MEA well medium was fully exchanged for Brainphys medium excluding BDNF, GDNF, and db-cAMP. The following day, medium was again fully replaced with medium excluding neurotrophic factors. After two hours, baseline recordings were performed. cLTP induction was performed by the addition of cAMP second messenger pathway modulators forskolin (LC Laboratories cat. #F-9929) and rolipram (Tocris cat. #0905). Medium containing 10x concentrations of forskolin and rolipram were added to MEA wells to give final concentrations of forskolin (50 µM) and rolipram (0.5µM). Wells treated with DMSO vehicle were subjected to identical medium changes as cLTP wells. 30 minutes after cLTP induction, MEA plates were recorded. Following the 30 minute time point, wells were washed with three full medium changes with Brainphys medium without BDNF, GDNF, and db-cAMP. Recordings were performed 4, 24, 48, and 72 hours post cLTP induction. Two-thirds of MEA well medium was removed and replaced following the 48 hour time point recording. cLTP MEA recordings were processed as described above. Due to large changes in intensity and frequency of network bursts, all recordings were reviewed to ensure accurate detection of network bursts. In some cases, manual thresholding of the spike histograms was performed to accurately detect high frequency and low intensity network bursting.

### Measurements of secreted Aβ^1-40^ and Aβ^1-42^

Concentrations of secreted Aβ^1-40^ and Aβ^1-42^ were measured from conditioned medium collected from 48-well MEA culture plates. Neurons were passaged to MEA plates and maintained as described above. Medium was conditioned for 96 hours prior to collection and approximately 150 µl was harvested. MEA electrical activity was recorded and monitored for neuronal firing and network burst development. DPP 14 timepoint was used for preburst Aβ measurements and DPP 35 was used for postbursting Aβ analysis. Secreted Aβ levels were measured using an MSD Aβ V-PLEX ELISA assay (MesoScale Discovery #151200E-2). Two technical replicates per MEA well were measured and final Aβ^1-40^ and Aβ^1-42^ concentrations were reported as the average of the two replicates.

### Statistical testing and plotting

Statistical testing was conducted in GraphPad Prism software (GraphPad Software). Dataset normality was determined with the D’Agostino and Pearson test. Parametric data were tested using two-tailed unpaired t-tests, paired t-tests, one-way ANOVA tests, or two-way ANOVA tests. Non-parametric data were analyzed with the Kolmogorov-Smirnov test. Tukey’s multiple comparisons testing was performed following one-way ANOVA testing and Sidak’s multiple comparisons testing was performed following two-way ANOVA testing. Plots were generated in R statistical software or GraphPad Prism. Data are presented as mean ± standard deviation (SD) or standard error of the mean (SEM), as appropriate. Representative immunocytochemistry images of analyzed neurons were generated in Imaris and signal brightness levels were optimized for signal visibility. MEA spike time histogram and raster plot traces were generated using Matlab.

## Results

### Widespread disruption of protein interactions among AMPA receptors, NMDA receptors, scaffolds and effectors

Based on our previous observation of reduced cell surface GLUR1 in *SORL1* KO neurons, we hypothesized that altered endosomal recycling due to *SORL1* deficiency alters the composition of synaptic proteins in excitatory cortical neurons. After bursting activity was confirmed in parallel cultures by MEA, approximately day post plating (DPP) 35 cultures were lysed and subjected to Quantitative Multiplex co-Immunoprecipitation (QMI), a mesoscale proteomics assay that reports the relative abundance of ∼400 binary interactions among 21 proteins expressed at the excitatory glutamate synapse (Brown 2019, Lautz 2018, **Figure 1A**, see **Table S1** for proteins and antibodies). Principal component analysis of QMI data matrices showed clear separation of wild-type and *SORL1* KO neurons across principal component 1, with additional effects of both batch (two separate differentiations) and cell line (two cell lines in duplicate per differentiation) (**Fig 1B**). To identify the specific interactions responsible for this effect, QMI data were processed using two independent statistical approaches: (1) adaptive nonparametric test with adjustable alpha cutoff (ANC) was designed specifically for QMI datasets (Smith 2016) and compares bead distributions for each of 400 interactions while correcting for multiple comparisons; while (2) correlation network analysis (CNA) (Horvath and Langfelder 2008) clusters interactions by similar behavior across all experimental samples, and asks if any co-regulated clusters, termed *modules*, correlate with experimental variables (**Fig 1A**). Only interactions that meet both CNA and ANC significance thresholds are reported hereafter.

CNA identified four modules that correlated with genotype, arbitrarily color-coded Blue (correlation coefficient (CC) = 0.86, p = 2 × 10^−5^), Magenta ( CC = 0.67, p = 0.005), Turquoise (CC=0.57, p = 0.02) and Black CC = 0.56, p = 0.03). Note, however, that Magenta and Black also correlated strongly with a Batch effect (CC = 0.93 p = 3 × 10^−7^ and CC = 0.8 p = 2 × 10^−4^, respectively) and Turquoise correlated with a Batch x Cell line effect (CC = 0.94 p = 4 × 10^−7^). Blue was the only module not confounded by batch and/or cell line. A heatmap of interactions that were ANC-significant for either batch 1 or 2, and a member of the four genotype-significant CNA modules (**Fig 1C**) demonstrates these effects. Interactions that were ANC-significant across both batches are in bold/italic. Immunoprecipitation target GLUR1, probe GLUR1 (abbreviated GLUR1_GLUR1) was the interaction most significantly correlated to the blue module, and was consistently reduced in *SORL1* KO cells across both batches (**Fig 1D**).

Interactions in which the IP and probe are for the same protein can indicate changes in abundance, multimerization, or changes in other co-associated proteins that may occlude antibody binding sites (Heavner 2021); prior published data showing reduced protein localization of GLUR1 supports the former interpretation (Mishra 2022). GLUR2_GLUR2 was significantly correlated to the black module, and was only significant by ANC for batch 2 (**Fig 1E**). These changes, combined with changes in GLUR1_GLUR2 in batch 2, suggest a shift in AMPA receptor subunit usage due to known endosomal GLUR1 trafficking deficits in *SORL1* mutants (Mishra 2022). In addition, alterations in Shank1_panShank (**Fig 1F**) or Homer_GLUR2 (**Fig 1G**) indicate changes to the Homer-Shank scaffolding network that supports AMPA receptor localization. Extensive changes to downstream effector molecules such as the RAS-GTPase SynGAP (exemplified by NMDAR2A_SynGAP, **Fig 1H**) or the SRC-family kinase FYN (**Fig 1I**) indicate altered signal transduction downstream of synaptic activity that could affect glutamate receptor properties or synaptic plasticity. A node-edge diagram limited to interactions that were ANC-CNA significant in both batches demonstrates the synapse-wide scope of the changes observed (**Fig 1J**). These data supported further investigations into synaptic activity in *SORL1* KO neurons.

### Loss of *SORL1* alters synapse morphology

QMI analysis demonstrated changes in multiple categories of excitatory synaptic proteins and their interactions. To determine if these changes altered the number and morphology of synapses we performed an immunocytochemical analysis of pre and post synaptic proteins. Wild-type and *SORL1* KO neuronal cultures were matured, fixed on DPP 30, and stained for SYNAPSIN and HOMER1 to identify pre and post synaptic structures, respectively. We performed three-dimensional analysis of pre and post synaptic marker signals. We found clear expression of SYNAPSIN and HOMER1 in the MAP2-positive somatodendritic compartments of wild-type and *SORL1* KO neurons (Fig 2A). Colocalization of the pre and postsynaptic markers shows that both neuronal genotypes form complete synaptic structures (Fig 2A, arrows). Several differences were detected in *SORL1* KO neurons. Consistent with QMI analysis, we detected increased expression of postsynaptic scaffolding protein HOMER1. *SORL1* KO HOMER1 puncta were larger and showed a greater sum fluorescence intensity (Fig 2B).

**Figure 2:**
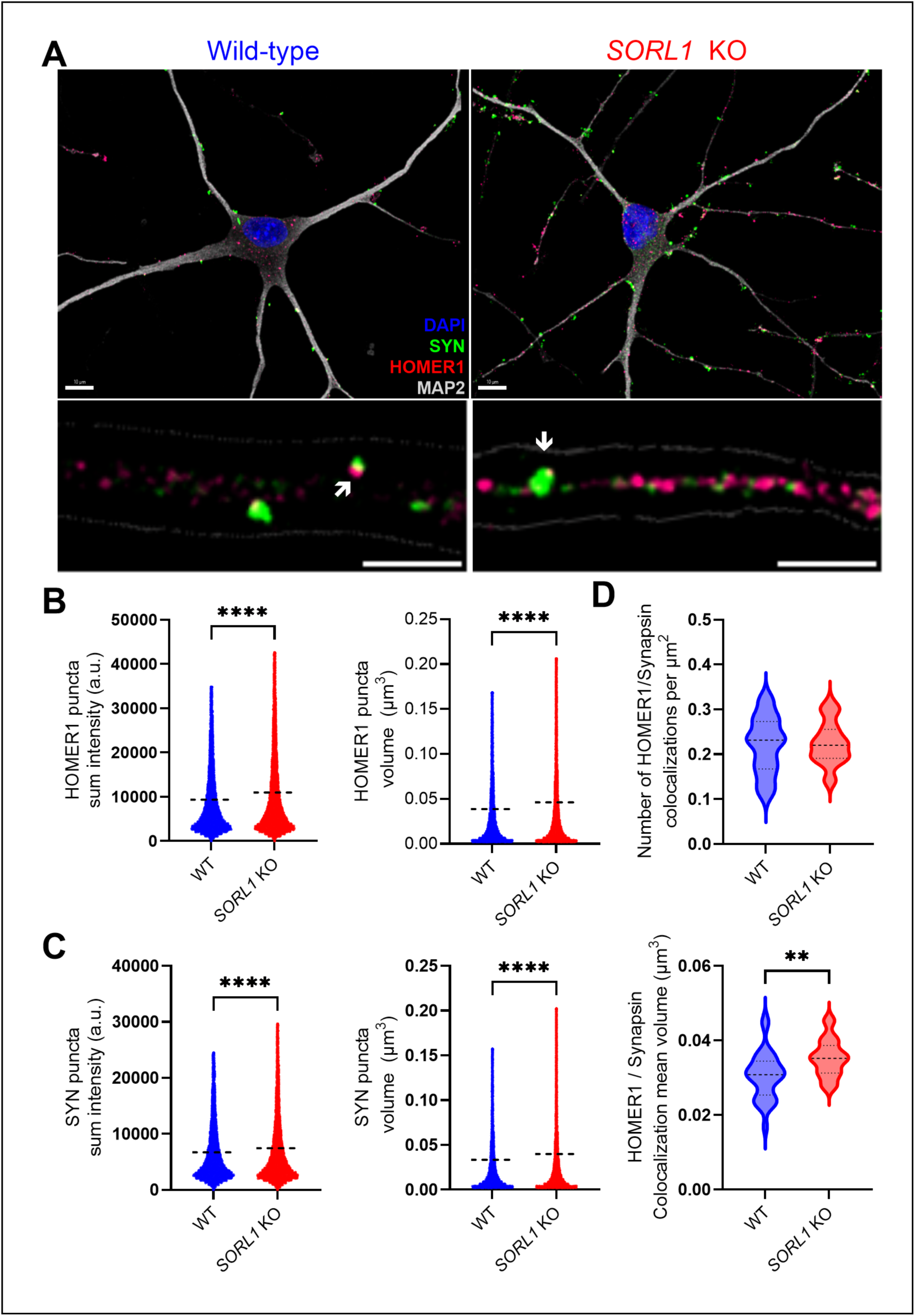
*SORL1 KO* neurons express enlarged pre and post synaptic densities. A: SORL1 deficient neurons exhibited larger and brighter pre and post synaptic structures. Representative micrographs of DPP 30 wild-type and SORL1 KO neurons, immunostained for pre and post synaptic markers SYNAPSIN (SYN) and HOMER1, and neuronal somatodendritic compartment marker MAP2. Lower panels: Higher magnification images of MAP2 positive dendrites. MAP2 signal borders indicated by grey outline. HOMER1 and SYNAPSIN colocalization highlighted with white arrow. B: Quantification of sum fluorescence intensity and volume of three-dimensional HOMER1 postsynaptic puncta. HOMER1 puncta in SORL1 KO neurons contained greater fluorescence signal and were larger than in isogenic wild-type control neurons. C: Quantification of sum fluorescence intensity and volume of three-dimensional SYNAPSIN (SYN) presynaptic puncta. SYNAPSIN puncta in SORL1 KO neurons contained greater fluorescence signal and were larger than in isogenic wild-type control neurons. D: Quantification of count and volume of three-dimensional puncta of HOMER1 and SYN colocalizations. No difference in the total number of colocalization puncta were detected between wild-type and SORL1 KO neurons, indicating similar levels of synapses. The volumes of colocalization puncta were larger in SORL1 KO neurons, suggesting larger synapses. Number of colocalizations was normalized to the total surface area of MAP2 volumes to control for image to image differences in number of neurons and dendrites. 80,000 to 140,000 HOMER1 puncta were analyzed and 40,000-60,000 SYNAPSIN puncta were analyzed from N = 32-34 images per genotype. Two clones per genotype were analyzed. HOMER1 and SYNAPSIN puncta quantification data points are plotted, with mean value indicated by horizontal dashed line. Colocalization metrics were calculated per cell and represented as violin plots with data median indicated by bolt dashed line and upper and lower quartiles represented by dotted lines. HOMER1 and SYNAPSIN puncta were determined to be non-normal (D’Agostino and Pearson test) analyzed for statistical significance with the Kolmogorov-Smirnov test. Outliers were identified using the ROUT (Q = 1%) method and removed for statistical testing and plotting. Statistical testing for colocalization quantifications were unpaired t-tests. Significance marked as follows: P < 0.01 (**), P < 0.0001 (****). Not significant comparisons are unmarked.

Staining of the presynaptic marker SYNAPSIN (SYN) revealed an increased fluorescence intensity and puncta volume in *SORL1* KO (Fig 2C), an analogous increase in the presynaptic compartment alongside the postsynaptic structure. Three-dimensional colocalization analysis revealed no difference in the number of colocalized HOMER1 and SYNAPSIN positive puncta, indicating that the number of synapses does not differ between *SORL1* KO neurons and isogenic controls (Fig 2D, upper). The volume of HOMER1 and SYNAPSIN colocalizations were larger in *SORL1* KO neurons, suggesting larger synapses (Fig 2D, lower). Taken together, these data indicate in *SORL1* KO neuronal synapses, altered synaptic structure is found on both sides of the synapse. We next compared pre and post synaptic protein signal in the somatic and dendritic compartments (Fig S1). The observed increased HOMER1 puncta volume and intensity in the whole-cell analysis was found in both the somatic and dendritic compartments of *SORL1* KO neurons (Fig S1 A,C). A small but significant decrease in SYNAPSIN puncta volume and intensity were found in the soma of *SORL1* deficient neurons (Fig S1 B). Conversely, SYNAPSIN puncta in the dendrites of *SORL1* KO neurons exhibited greater intensity and volume than isogenic control neurons (Fig S1 D), demonstrating that the observed differences in presynaptic SYNAPSIN staining we found in the whole-cell analysis (Fig 2) was primarily driven by differences in the *SORL1* KO dendrites. *SORL1* KO neurons show clear differences in excitatory synaptic densities, recapitulating the increased HOMER1 interactions identified by QMI (Fig 1) and accompanying changes in the presynaptic compartment.

### *SORL1* KO neurons have shifted AMPA receptor subunit expression

We have shown previously that *SORL1* KO neurons have impaired trafficking of the glutamate AMPA receptor subunit GLUR1 (Mishra 2022) and here we find altered interactions of GLUR1 and GLUR2 subunits in *SORL1* KO neurons with QMI analysis. We performed an immunocytochemical assay of GLUR1 and GLUR2 expression in matured wild-type and *SORL1* KO neurons to further examine the expression profiles of the AMPA receptor subunits. We found GLUR1 and GLUR2 expression in wild-type and *SORL1* KO neurons (Fig 3A). GLUR1 puncta in wild-type neurons has on average greater sum fluorescence intensities and volumes compared to those in *SORL1* KO neurons (Fig 3B). Conversely, *SORL1* KO neurons had, on average, GLUR2 puncta with greater sum fluorescence intensities and volumes than isogenic wild-type control neurons (Fig 3C). These patterns of expression align with GLUR1 and GLUR2 interaction changes observed in QMI (FIGURE 1) and reflect a shift in the balance of GLUR1 and GLUR2 subunit expression profiles when SORL1 dependent trafficking is disrupted. We next measured GLUR1 and GLUR2 puncta in somatic and dendritic compartments and found little difference in comparison to whole-cell analysis (Fig S2), other than GLUR1 puncta volume being similar in the dendrites of wild-type and *SORL1* KO neurons. These data show that differences in AMPA receptor subunit expression are occurring cell-wide and not differently in somatic and dendritic compartments. These results add to our previous findings of GLUR1 trafficking being disrupted.

**Figure 3:**
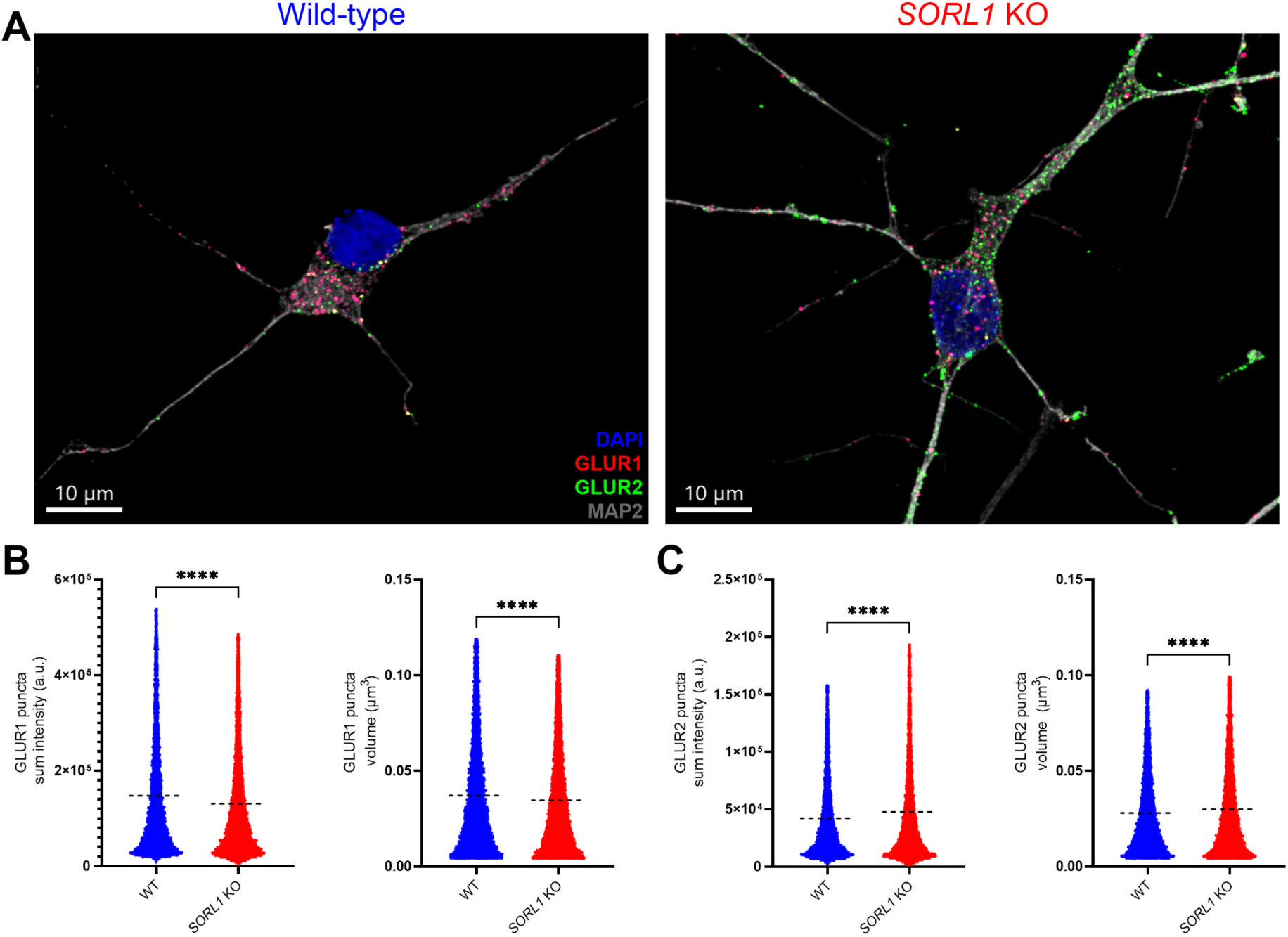
*SORL1* deficient neurons exhibit abnormal AMPA subunit GLUR1 and GLUR2 levels. A: Representative images of isogenic wild-type and *SORL1* KO neurons, immunostained for AMPA receptor subunits GluR1, GluR2, somatodendritic neuronal marker MAP2, and DAPI. B: Quantification of GluR1 puncta sum fluorescence intensity and volume from wild-type and *SORL1* KO neurons. GluR1 puncta in *SORL1* KO neurons were smaller and had lower sum fluorescence intensity. C: GluR2 puncta sum fluorescence intensity and volumes from wild-type and *SORL1* KO neurons. Immunostained GluR2 puncta were brighter and larger in volume in *SORL1* KO neurons compared to wild-type neurons. N = 10,000-12,000 GluR1 and GluR2 puncta were analyzed from 32-34 images per genotype. Two clones per genotype were analyzed. ICC puncta data points plotted with mean values indicated by horizontal dashed line. Outliers were identified using the ROUT (Q = 1%) method and removed for statistical testing and plotting. Data were tested for normality and were determined to be not normally distributed (D’Agostino and Pearson test). Statistical testing for all quantifications was the Kolmogorov-

### *SORL1* deficient neurons are hyperexcitable

Broad changes in synaptic protein composition suggest physiological consequences of loss of *SORL1.* Laboratory models of AD repeatedly demonstrate hyperexcitable neuronal phenotypes at the network and cellular level (Busche 2008, Ghatak 2019, Lerdkrai 2018, Targa Dias Anastacio 2022). Here, to assess how *SORL1* contributes to neuronal function, we measured the spiking activity of *SORL1* KO and isogenic control hiPSC derived neurons on multiple electrode array (MEA) plates for sixty days post differentiation. Over this time period *SORL1* KO cultures exhibited several hyperexcitable phenotypes (Fig 4). *SORL1* KO neurons generated more action potentials than isogenic controls, with *SORL1* KO cultures generating greater mean firing rates (Fig 4B). The difference in firing rate was larger in more mature cultures. The timing of KO neuron’s spiking was also less regular, as demonstrated by the higher coefficient of variation of the inter-spike interval in *SORL1* KO neuron cultures (Fig 4C). Similarly, this increased variability in timing in *SORL1* KO neurons was greatest in more mature cultures.

**Figure 4:**
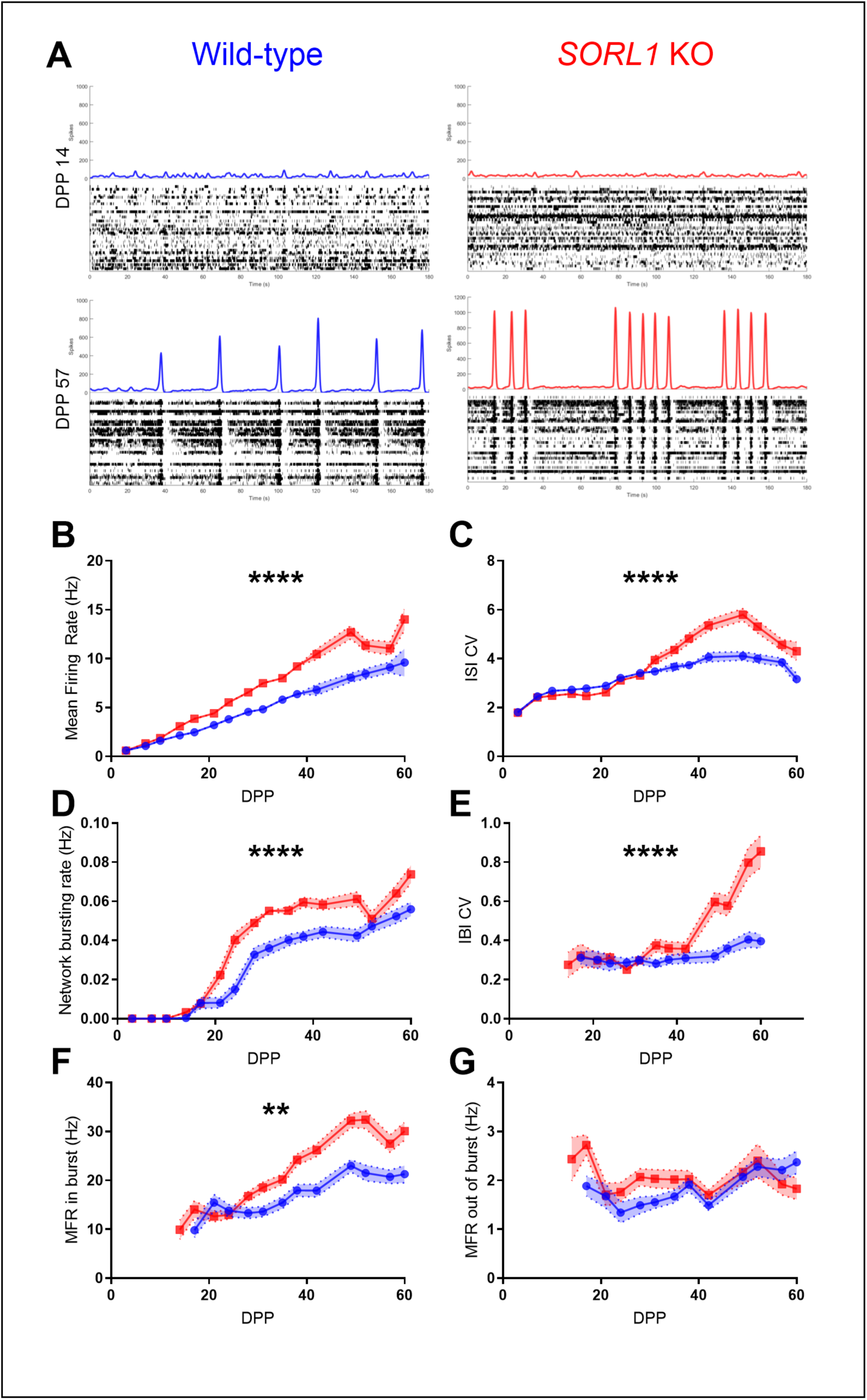
*SORL1* KO neurons are hyperexcitable. A: Representative spike histograms (upper panel) and raster plots (lower panel) of wild-type and *SORL1* KO neuronal cultures plated on multi-electrode arrays (MEAs) at 14 and 57 DPP, before and after the onset of synchronized network bursting. B: Time course of mean firing rates of wild-type and *SORL1* KO neuron cultures. C: Time course of the mean coefficient of variation (CV) of inter-spike intervals (ISI) of action potentials generated by wild-type and *SORL1* KO neuron cultures. D: Time course of the mean rate of network bursts generated by wild-type and *SORL1* KO cultures. E: Time course of the mean coefficient of variation of inter-burst intervals (IBI) of network bursts generated by wild-type and *SORL1* KO neuron cultures. *SORL1* KO neurons generated more spikes with more variable timing and this pattern was conserved at the network activity level, reflected by the differences between wild-type and *SORL1* KO neuron culture network bursting activity. F: Time course of the mean firing rate of spikes generated during a network burst. G: Time course of the mean firing rate of spikes generated outside of a network burst. Neuronal firing rates were higher during *SORL1* KO neuron network bursts while no difference was detected in the network burst interstices. N = 54-63 MEA wells analyzed per genotype. Two clones per genotype were analyzed. Data represented as MEAN ± SEM. Two-way ANOVA statistical testing was used for all quantifications. Significance of the interaction of genotype and time marked on the plots as follows: P < 0.05 (*), P < 0.01 (**), P < 0.001 (***), P < 0.0001 (****). Unsignificant comparisons are unmarked.

After approximately three weeks, KO and control neuronal cultures began to exhibit coordinated activity indicative of neurotransmission-driven network firing, or network bursting, consistent with pre and post synaptic protein staining (Fig 4A, Fig 2). *SORL1* KO neuronal hyperexcitability was conserved at the network level and KO cultures generated more network bursts compared to isogenic controls (Fig 4D). *SORL1* KO culture network bursting also showed more irregularity in the timing of the interburst interval (IBI), reflected by an increased coefficient of variation in the inter-burst interval (Fig 4E). The hyperexcitability of *SORL1* KO cultures was accelerated by the onset of network bursting, so we examined the rates of spiking inside and outside the network burst window. Surprisingly, we found no difference between genotypes in the mean firing rate outside of coordinated bursting (Fig 4G). Within the duration of the burst, however, *SORL1* KO neurons fired at an increased rate (Fig 4F). These results indicate that *SORL1* KO neuron hyperexcitability is driven during coordinated network bursts.

### Hyperexcitability in *SORL1* deficient neurons is glutamate dependent

*In vitro* cultures of hiPSC excitatory neurons developing coordinated network bursting is an established phenomenon (Heikkila 2009, Plumbly 2019). The hyperactivity of *SORL1* KO network activity and changes observed in excitatory synapse protein interactions and morphology implicate changes to synapses could be driving *SORL1* KO neuron hyperexcitability. Here, we tested if *SORL1* KO hyperexcitability persisted with blockade of fast ionotropic glutamatergic AMPA and NMDA receptors. Following the development of network bursting, wild-type and *SORL1* KO neuronal cultures were treated with 10 µM NBQX and 50 µM APV to block AMPA and NMDA mediated neurotransmission. Network bursting was completely abolished in both wild-type and *SORL1* KO cultures with treatment (Fig 5A). Mean firing rates and coefficient of variation of inter-spike intervals also decreased with antagonist application (Fig 5B,C). Notably, the differences in these metrics between wild-type and *SORL1* KO neurons were eliminated when glutamate neurotransmission was blocked and restored after antagonist washout. Inhibitory neurotransmission shapes network activity in other models (Odawara 2016, Plumbly 2019, Cobb 1995) and dual-SMAD neuronal differentiation can lead to the expression of inhibitory neuron genes and proteins (Israel 2012). We found no significant contribution of GABA_A_ mediated inhibitory neurotransmission in our neural networks. Application of 10 µM gabazine leads to small, non-significant increases in firing rates of wild-type and *SORL1* KO neuronal cultures (Fig 5D, Fig S3). No changes to network bursting rate were observed (Fig S3B) and the coefficient of variation of the inter-burst interval (Fig S3C) with gabazine treatment. Wild-type and *SORL1* KO neurons showed similar levels and patterns of spiking when glutamate receptors were blocked, suggesting that the hyperexcitability phenotypes we found are the result of abnormal excitatory glutamate neurotransmission.

**Figure 5:**
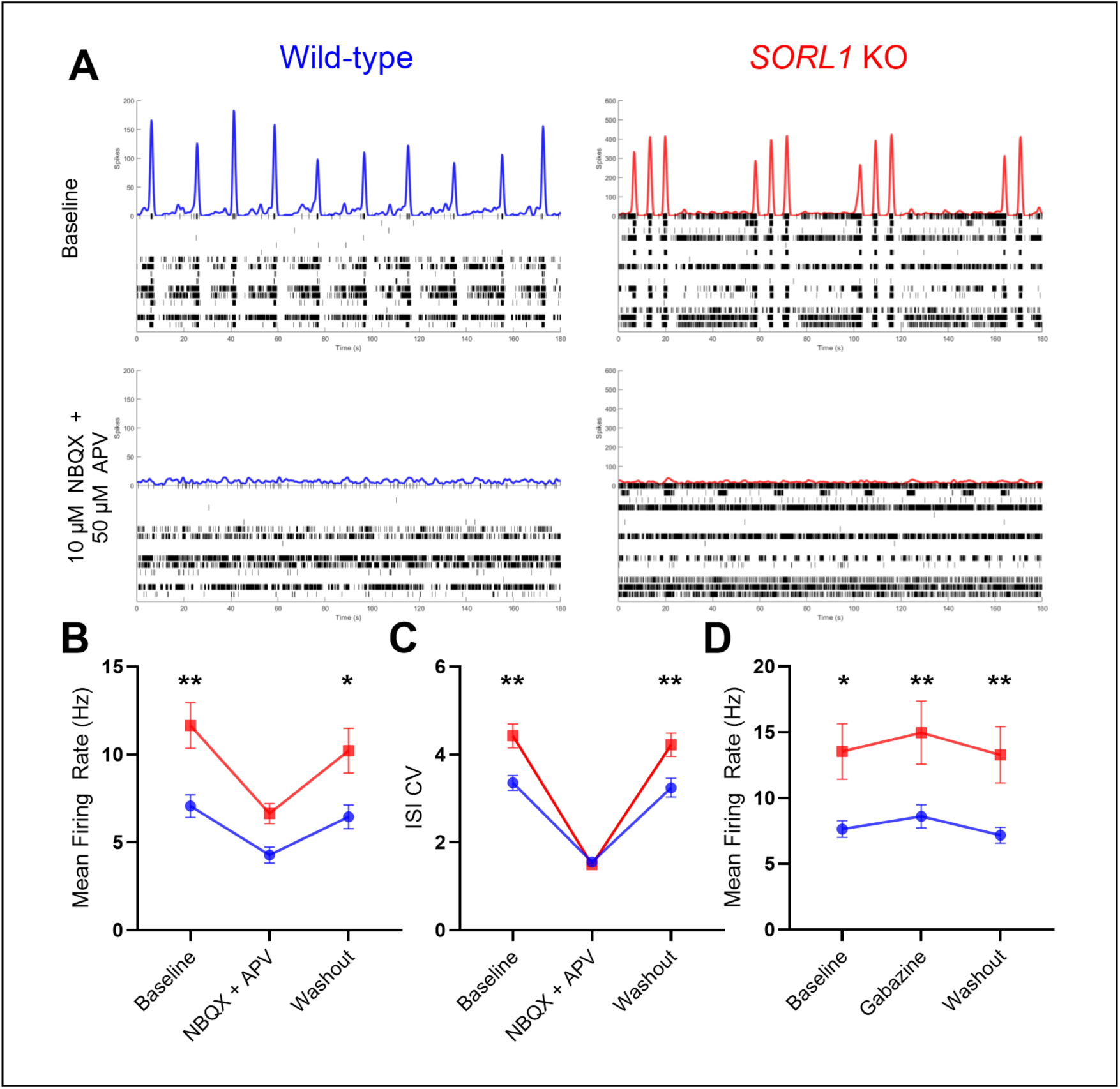
*SORL1* KO neuron hyperexcitability is driven by glutamatergic neurotransmission. A: Representative *SORL1* KO neuron culture spike histogram (upper panel) and raster plot (lower panel) before and after treatment with glutamate receptor AMPA and NMDA antagonists NBQX and APV (10 μM and 50 μM respectively). Glutamate receptor blockade eliminates network bursting, indicating that network activity is driven primarily by glutamatergic neurotransmission. B: Mean firing rates of wild-type and *SORL1* KO cultures treated with NBQX and APV. C: Coefficient of variation of the inter-spike interval of wild-type and *SORL1* KO neuron cultures treated with NBQX and APV. Wild-type and *SORL1* KO neurons have significantly different mean firing rates and CVs of ISIs in baseline and washout conditions but are similar when glutamate neurotransmission is blocked, demonstrating that *SORL1* hyperactivity is driven by glutamate receptor activity. D: Mean firing rate of wild-type and *SORL1* KO neurons treated with the GABA_A_ receptor antagonist Gabazine (10 μM). A small, non-significant increase in firing rate was measured with Gabazine treatment in both wild-type and *SORL1* KO neurons, showing that inhibitory GABAergic neurotransmission does not play a role in *SORL1* KO hyperactivity. N = 24 wild-type cultures and 36 *SORL1* KO cultures were used for NBQX and APV experiments at approximately DPP 45. Two clones per genotype were analyzed for NBQX and APV experiments. N = 27 wild-type and *SORL1* KO cultures were used for Gabazine experiments at approximately DPP 45. Two wild-type clones and one *SORL1* KO clone were analyzed for Gabazine experiments. Data represented as MEAN ± SEM. Two-way ANOVA statistical testing was performed for all quantifications. P-values of Sidak’s multiple comparisons test between wild-type and *SORL1* KO genotypes at baseline, antagonist condition, and washout are represented on the plots with significance values marked as: P ≤ 0.05 (*), P ≤ 0.01 (**). Non-significant comparisons are unmarked.

### Hyperexcitability in *SORL1* KO neurons is not solely due to increases in secreted amyloid beta

*SORL1* KO and *SORL1* loss-of-function variant hiPSC neurons secrete more soluble Aβ as a result of impaired trafficking (Knupp 2020, Mishra 2023). Soluble Aβ causes increased neuronal activity and impaired synaptic plasticity in hiPSC neurons and other laboratory models (Zott 2019, Ping 2015, Busche 2008, Palop 2010). To examine if increased Aβ could explain the physiological phenotypes we observe in *SORL1* KO neurons, we compared neurophysiological activity of neural cultures using MEA and harvested medium from MEA wells to measure secreted Aβ protein concentrations of *SORL1* KO neurons, an isogenic line carrying homozygous amyloid precursor protein (APP) mutations KM670 and 671NL, the APP Swedish mutations (APP^Swe+/+^), and an isogenic wild-type control. APP^Swe+/+^ neurons secreted much higher levels of Aβ 1-40 and 1-42 than wild-type control and *SORL1* KO neurons (Fig 6A-D) (Shin and Evitts, 2023). Consistent with prior studies, APP^Swe+/+^ neurons were hyperactive compared to isogenic wild-type controls (Fig 6E,F). APP^Swe+/+^ neurons successfully developed coordinated network bursting, demonstrating that the increased levels of soluble Aβ does not interfere with network synapse development. Interestingly, mean firing rates of APP^Swe+/+^ neurons were similar to *SORL1* KO neurons prior to the development of network bursting (Fig 6E). After network bursting was established, APP^Swe+/+^ neurons continued to fire more than isogenic controls but significantly less than *SORL1* KO neurons (Fig 6F). All cell lines generated more Aβ 1-42 after developing bursting and wild-type and *SORL1* KO neurons also secreted significantly more Aβ 1-40 (Fig S4). Network bursting *SORL1* KO neurons generated significantly more spiking than APP^Swe+/+^ neurons (Fig 6F). Despite the increased level of neuronal spiking observed prior to and after network bursting in APP^Swe+/+^ neurons compared with control neurons, APP^Swe+/+^ neurons did not generate more network bursts compared with WT neuronal cultures, in contrast with *SORL1* KO cultures, which maintained their hyperactivity at the network level (Fig 6G). These results emphasize the *SORL1* KO driven alterations to excitatory synapse composition being the principal driver of *SORL1* KO hyperexcitability. When firing rates and concentrations of secreted Aβ are compared directly, we found that the hyperexcitability observed in *SORL1* KO neuronal cultures is not due solely to increased Aβ secretion (Fig 6H).

**Figure 6:**
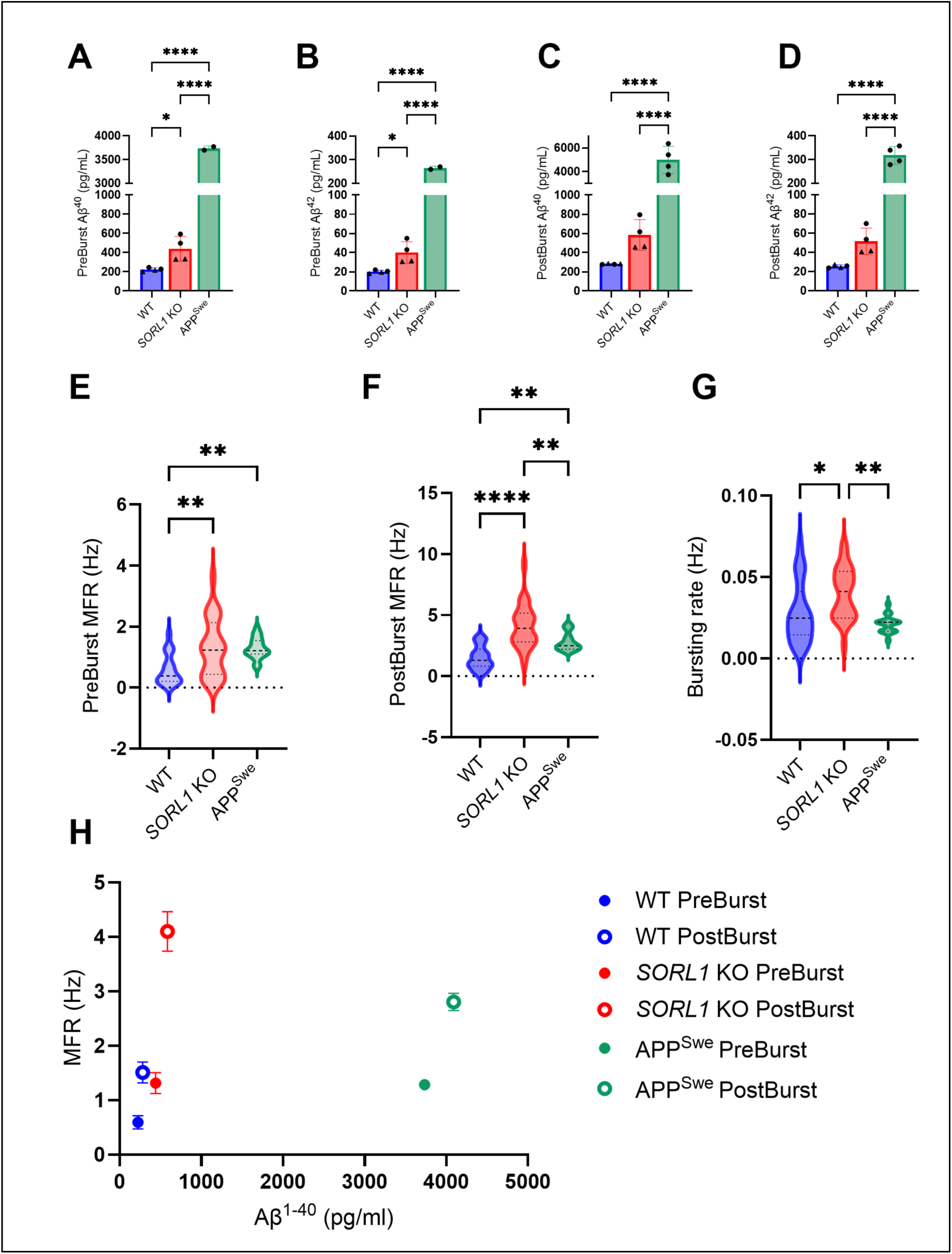
*SORL1* KO neuron hyperexcitability is independent of secreted amyloid beta. A-B: Concentrations of secreted Aβ^1-40^ and Aβ^1-42^ measured from conditioned medium collected from DPP 14 MEA cultures of wild-type, *SORL1* KO, and APP^Swe +/+^ neurons, before network bursting has been established. C-D: Concentrations of secreted Aβ^1-40^ and Aβ^1-42^ peptide measured from conditioned medium collected from DPP 35 MEA cultures of wild-type, *SORL1* KO, and APP^Swe +/+^ neurons, after network bursting has been begun. At both timepoints, prebursting (DPP 14) and postbursting (DPP 30), APP^Swe +/+^ neurons secrete significantly greater levels of Aβ^1-40^ and Aβ^1-42^ than isogenic wild-type controls and *SORL1* KO neurons. N = 4 wild-type and *SORL1* KO cultures. N = 2 APP^Swe +/+^ cultures. 2 clones of wild-type and *SORL1* KO cell lines were used, 1 clone of APP^Swe +/+^ was used. Each point represents the average of two technical replicates measured from conditioned medium collected from one MEA well ± SD. One-way ANOVAs were used for statistical testing. Tukey’s multiple comparisons test results are represented on the plot with significance marked: E-F: Mean firing rates or wild-type, *SORL1* KO, and APP^Swe +/+^ neurons at conditioned medium collection timepoints ,DPP 14 and DPP 30. *SORL1* KO and APP^Swe +/+^ neurons exhibited increased firing rates compared to isogenic wild-type controls in prebursting conditions. After the establishment of network bursting, *SORL1* KO neurons generated action potentials at a greater rate than wild-type and APP^Swe +/+^ neurons. G: Network bursting rates of wild-type, *SORL1* KO, and APP^Swe +/+^ at DPP 30. *SORL1* KO cultures generated network bursts at a greater rate than both wild-type and APP^Swe +/+^ cultures, demonstrating that network hyperactivity is not solely dependent on secreted levels of Aβ^1-40^ or Aβ^1-42^. N = wild-type (24), *SORL1* KO (24), and APP^Swe +/+^(20). Data represented as violin plots with data median indicated by bolt dashed line and upper and lower quartiles represented by dotted lines. One-way ANOVAs were used for statistical testing. Tukey’s multiple comparisons test results are represented on the plot with significance marked: H: Scatter plot of secreted Aβ^1-40^ peptide concentration vs mean firing rate for both prebursting and postbursting timepoints. Despite the greatly increased levels of Aβ^1-40^ and Aβ^1-42^ peptides present in the medium, the mean firing rate and network bursting rate of APP^Swe +/+^ cultures fail to recapitulate the levels of hyperexcitability observed in *SORL1* KO neurons. Data points represent the mean values of each genotype’s Aβ^1-40^ peptide concentration and mean firing rate ± SEM. Significance indicated by: P ≤ 0.05 (*), P ≤ 0.01 (**), P ≤ 0.001 (***), and P ≤ 0.0001 (****). Not significant comparisons are not marked.

### Loss of *SORL1* inhibits long-term potentiation induction

Long-term potentiation (LTP) is a form of synaptic plasticity in which GLUR1-containing AMPA receptors are rapidly trafficked to the postsynaptic membrane in response to high levels of presynaptic activity (Malenka 2004). Inhibited LTP has been linked to AD in several models, and the SORLA binding partner retromer is a trafficking complex that is necessary for LTP (Temkin 2017, Simoes 2021) Chemical LTP (cLTP) protocols are effective approaches to inducing similar changes in neurons, and are especially useful *in vitro* where networks lack organized afferent inputs. The cAMP second messenger pathway can be positively modulated to induced LTP in hiPSC neuronal cultures. Network bursting is dependent upon glutamate receptor activity (Fig.5) and induction of cAMP cLTP results in increased levels of network bursting in hiPSC derived dopaminergic neuron networks (Pre 2022). Here, we utilized a cLTP protocol developed in Pre et al 2022 to examine the network activity of *SORL1* KO neuronal cultures that have undergone LTP induction. The adenylyl cyclase activator forskolin was used in combination with the phosphodiesterase inhibitor rolipram to stimulate the cAMP second messenger pathway and prevent cAMP degradation. We induced cLTP in wild-type and *SORL1* KO neurons plated on MEA culture plates and assayed the electrical activity for 72 hours post induction. Wild-type neurons displayed increases in network bursting rate that lasted up to 72 hours post induction (Fig 7A,B). In contrast, *SORL1* KO neurons failed to recover network bursting activity for up to 24 hours post induction (Fig 7 A,B), with significantly lower fold change bursting rates in comparison with wild-type cultures. Mean firing rates measured from MEA cultures increased in wild-type cultures slower than the increase in network bursting, similar to what has been described in dopaminergic neuronal network cLTP responses (Pre 2022). cLTP treated *SORL1* KO neurons showed no change in mean firing rates compared to pretreatment baseline conditions (Fig 7C). Interestingly, *SORL1* KO neurons do exhibit an increased network bursting rate once network activity has resumed and the differences between wild-type and *SORL1* KO network bursting fades at the 48 hour timepoint post induction (Fig 7A,B). These data demonstrate that *SORL1* is an important component of activity driven increases of excitatory synaptic strength.

**Figure 7:**
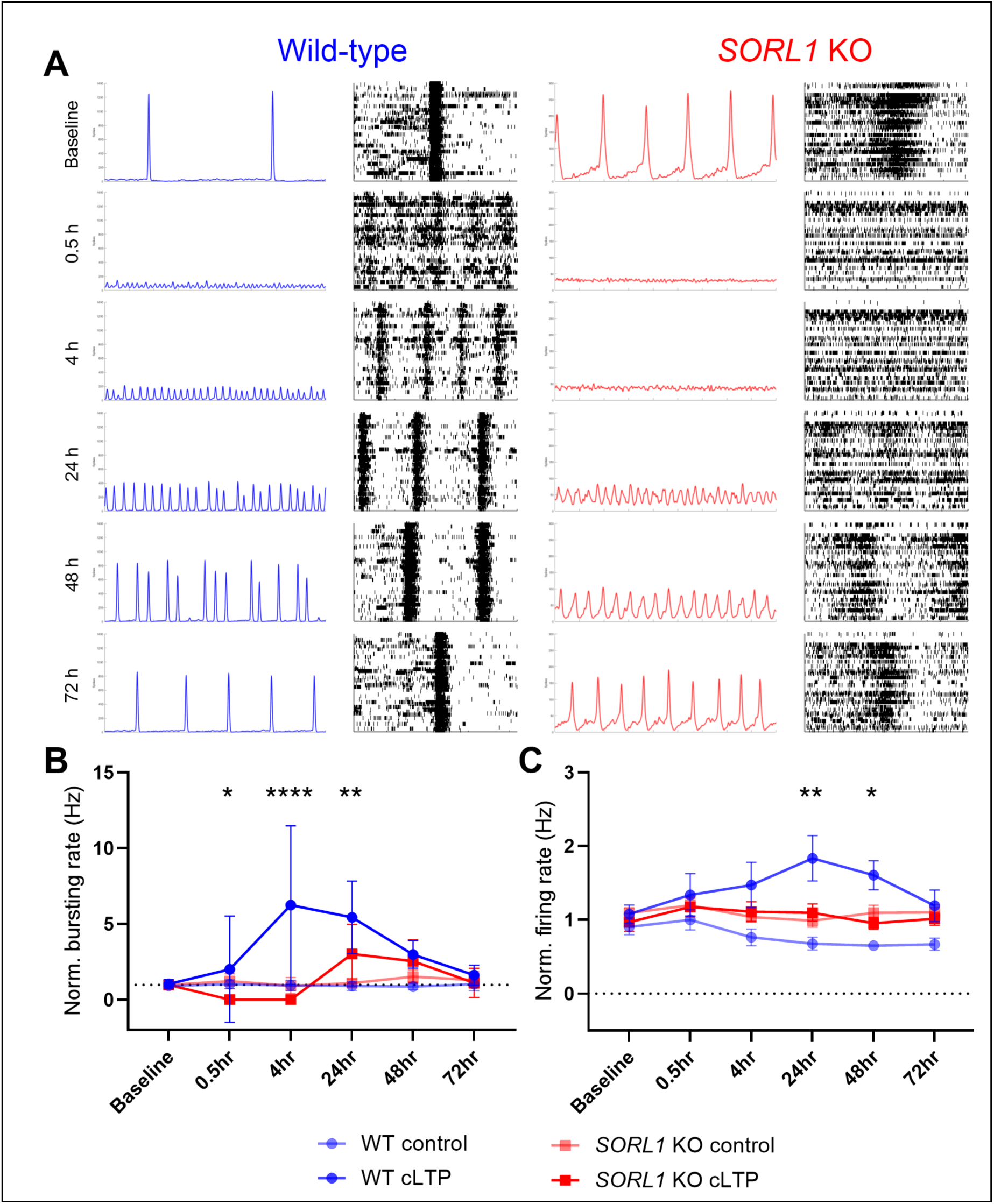
*SORL1* KO neurons exhibit inhibited long-term potentiation. A: Representative spike histograms (left panels) and raster plots (right panels) of wild-type and *SORL1* KO cultures across the time course of cLTP treatment and recovery. Raster plots represent twenty seconds of the full recording shown by the spike histogram, highlighting the pattern of activity that constitutes a network burst. Wild-type and *SORL1* KO neurons display a breakdown of synaptically-driven activity 30 minutes following chemical LTP (cLTP) induction utilizing forskolin and rolipram. B: Time course of fold change in network bursting rate, normalized to baseline values, of wild-type and *SORL1* KO neurons following cLTP induction. Network bursting in wild-type cultures resumed in wild-type cultures within four hours of cLTP induction but *SORL1* KO neurons failed to recover by this time. Network bursting in *SORL1* KO neurons is reestablished on average by 24 hours, albeit at a smaller relative rate compared with wild-type neurons. No change was observed with control vehicle treatment C: Time course of fold change in mean firing rate, normalized to baseline values, of wild-type and *SORL1* KO neurons following cLTP induction. *SORL1* KO neurons did not exhibit increased firing rates following cLTP treatment, in contrast with wild-type neurons. N = 9-11 MEA cultures of wild-type and *SORL1* KO neurons. 2 clones of each genotype were used for cLTP experiments. Two-way ANOVA and Tukey’s multiple comparisons statistical tests were performed. Significance between wild-type cLTP and *SORL1* KO cLTP groups are marked on B and C plots with P ≤ 0.05 (*), P ≤ 0.01 (**), P ≤ 0.001 (***), and P ≤ 0.0001 (****). Non-significant differences are not marked.

## Discussion

In this study, we report that loss of *SORL1* expression affects excitatory synapse composition, morphology, and neuronal function, including the impairment of long-term potentiation. Synaptic dysfunction is a very early event in AD progression (Pelucchi 2022) and the main contributor to cognitive decline (Terry 1991), thus preventing this impairment at an early stage would be a beneficial therapeutic option for AD.

The relationship between neuronal hyperexcitability with synaptic function and dysfunction is complex. In AD, hyperexcitability has been attributed to increases in Aβ molecules which can directly depolarize neurons and disrupt inhibitory circuits, and pTau pathology which can impair axonal transport and destabilize neuronal networks (Pelucchi 2022). However, impaired endosomal recycling has emerged as a hypothesis that is upstream of both Aβ and pTau, and can contribute to both of these pathologies (Small and Petsko 2015). Multiple proteins necessary for normal synaptic composition are trafficked through the endosomal network (Park 2004, Dong 2024). Based on our previous work (Mishra 2022), we hypothesized that impaired endosomal recycling in *SORL1* deficient neurons would alter synaptic protein trafficking. Using an unbiased approach, QMI, we show that multiple synaptic proteins, including receptors and scaffolding proteins, show altered protein-protein interactions.

HOMER1 and PSD95 postsynaptic scaffolding proteins both demonstrate increased interactions in *SORL1* KO neurons, suggesting post synaptic density alterations. This is corroborated by our immunostaining results showing larger volumes and intensities of HOMER1 signal (Fig 2). We also observed increased fluorescence intensity and volume of SYNAPSIN puncta. While these increases could explain the hyperexcitability we observe, combined with reduced long-term potentiation, this could represent an ultimately maladaptive response to altered synaptic protein trafficking by which the neuron attempts to maintain synaptic strength by increasing synaptic size and expanding the molecular machinery for synaptic transmission, such as the accumulation of scaffolding proteins. Interestingly, we found similar numbers of synapses in wild-type and *SORL1* KO neurons (Fig 2), showing that *SORL1* trafficking contributes to the maintenance of proteins at synapses but does not result in compensatory formation of new synapses. Similarly, the hyperactivity of *SORL1* deficient neurons does not cause a homeostatic compensatory decrease in synapses to normalize neuronal firing levels.

We also observe alterations in AMPA receptor subunits GLUR1 and GLUR2 both by QMI and by immunostaining. There is a significant decrease in GLUR1-GLUR1 interactions in *SORL1* KO neurons, which corroborates our previous findings of reduced GLUR1 subunits trafficked to the neuronal membrane. Loss of GLUR1 on neuronal membranes has also been seen in models of retromer deficiency, suggesting that SORLA-retromer trafficking is a key mechanism for maintaining function AMPA receptors. We observe increased interactions between GLUR2 and increased intensity in GLUR2 immunostaining. While GLUR2 is also trafficked via retromer (Tian 2015), its interaction with SORLA is not clear. In a mouse neuronal model of VPS35 deficiency, decreased synaptic transmission could be rescued by GLUR2 overexpression (Tian 2015).

Thus, increased GLUR2 levels in our model may be a compensatory effect. GLUR2 subunit incorporation into AMPA receptors prolongs the decay time of synaptic currents, and leads to more efficient EPSP-AP coupling (Savtchouk 2011), which could contribute to the increased bursting we observed. In addition, GLUR2 subunits control the calcium permeability of AMPA receptors and block calcium influx after post transcriptional Q/R site editing. Decreased GLUR2 editing in the neurodegenerative disease amyotrophic lateral sclerosis (ALS) contributes to excitotoxicity in motor neurons (Kwak 2010). Similar mechanisms may be occurring when SORLA mediated transport is disrupted and shifts AMPA receptor subunit levels, as we have demonstrated here.

Blocking glutamate neurotransmission and synchronized network bursting normalizes *SORL1* KO neuron hyperexcitability demonstrating that aberrant firing rates are driven by glutamate neurotransmission. Wild-type and *SORL1* KO neuron firing rates are similar outside of synchronized network bursts, further support for a functional consequence of the resulting excitatory synaptic protein shifts we observed. Larger synapses are correlated with stronger synaptic currents (Hazan 2020, Holler 2021) and we observe larger HOMER1-SYNAPSIN colocalization volumes which could contribute to the hyperactivity observed in our cells.

Changes in excitatory-inhibitory balance has been implicated in AD-driven hyperactivity but we find no effect of blocking GABA_A_ inhibitory receptors (Figs 5, S3) beyond a small, non-significant increase in firing rate in both wild-type and *SORL1* KO neurons, making impaired inhibition a less likely cause in this model. In our previous study (Mishra 2022), bulk RNA sequencing showed increases in ion channel expression, which may be contributing to hyperexcitability independently of changes at the synapse.

The hyperexcitability phenotypes we find here are consistent with Raska, Satkova, and Plesingrova et al (co-submitted with this manuscript). There, the authors find hyperactivity in not only *SORL1* KO neurons but also the missense variant p.Y1816C. These results build upon other reports of detrimental effects of *SORL1* pathogenic variants (Mishra 2023, Fazeli 2024-1, Fazeli 2024-2, Rovelet-Lecrux 2021) and demonstrate that neuronal function is dependent upon functional SORLA protein and not affected solely by its complete loss of protein expression. Additionally, their measurements are taken from a different parental cell line and generated utilizing a transcription factor induced differentiation protocol, distinct from the dual-SMAD inhibition protocol used here, emphasizing that the critical component in our studies to be missing or malfunctioning *SORL1*. Together, these results demonstrate the robustness of *SORL1*’s importance for healthy neuronal function and contributions to early disease pathology.

Our findings demonstrate that SORL1 deficiency disrupts synaptic homeostasis through a mechanism that is largely independent of Aβ. While increases in Aβ due to loss of *SORL1* are documented (Knupp 2020) and may contribute to some of the hyperexcitability we observe, neurons with the APP Swedish mutation secrete more than 5X the amount of Aβ and are less hyperactive after the onset of synchronized, synaptically driven network activity. This suggests that the primary driver of neuronal hyperexcitability in *SORL1* deficient neurons is due to altered synapse composition as a result of SORLA-dependent endosomal trafficking and recycling.

Synaptic plasticity is dependent on the movement of receptors and proteins at the synapse, and deficits in long-term potentiation have been demonstrated in AD models and brains (Walsh 2002, Di Lorenzo 2017) and SORLA’s binding partner retromer (Temkin 2017, Simoes 2021). Here, we report the novel finding that loss of *SORL1* inhibits the ability of hiPSC neurons to induce LTP in response to chemical stimulation. Recycling endosomes near the post synaptic structure hold and shuttle AMPA receptors for LTP induction (Park 2004) and *SORL1* deficient neurons demonstrate recycling endosome swelling and increased co-localization with GLUR1 AMPA subunits (Mishra 2022). The lowered levels of response to cLTP induction we see here are likely due to a failure in the movement of GLUR1-containing AMPA receptors in response to induction. Interestingly, we do see a slow response to cLTP induction in *SORL1* KO neurons, showing that synaptic plasticity isn’t completely lost, reflecting low levels of recycling endosome to post synaptic membrane trafficking or diffusion of extrasynaptic AMPA receptors.

## Conclusions

Our work supports a model where synaptic trafficking defects represent an early and potentially targetable mechanism in AD pathogenesis. These insights suggest that therapeutic interventions aimed at restoring proper endosomal function or correcting synaptic protein trafficking may offer effective strategies for preventing cognitive decline in AD.

## List of abbreviations

Aβ: Amyloid beta
AD: Alzheimer’s disease
ANC: Adaptive nonparametric test with adjustable alpha cutoff
APOE: Apolipoprotein E
APP: Amyloid precursor protein
BDNF: Brain-derived neurotrophic factor
cLTP: Chemical long-term potentiation
CNA: Correlation network analysis
DPP: Days post plating
ELN: Endo-lysosomal network
GDNF: Glial derived neurotrophic factor
GLUR1: Glutamate receptor 1
GLUR2: Glutamate receptor 2
GWAS: Genome-wide association study
hiPSC: Human induced pluripotent stem cell
KO: Knock-out
MEA: Multi-electrode array
pTau: hyper-phosphorylated tau protein
QMI: Quantitative multiplex immunoprecipitation
SORL1: Sortilin-related receptor 1 (gene)
SORLA: Sortilin-related receptor with A-type repeats (protein)

## Declarations

### Ethics approval

Not applicable.

### Consent for publication

Not applicable.

### Availability of data and materials

All data and material in the current study are available from the corresponding author on reasonable request.

### Author’s contributions

Conceptualization-C.A.W., J.E.Y. Methodology and Analysis: C.A.W, S.E.R, V.S, S.E.P.S. Writing-C.A.W., J.E.Y., Revising and Editing-S.E.R, S.E.P.S. Funding Acquisition-J.E.Y., C.A.W.

## Acknowledgements

We thank Drs. Olav Andersen, Scott Small and Greg Petsko for helpful discussions. We thank the members of the Young Laboratory and the UW Alzheimer’s research community for critical feedback and discussions on this work.

---

**Figure S1:**
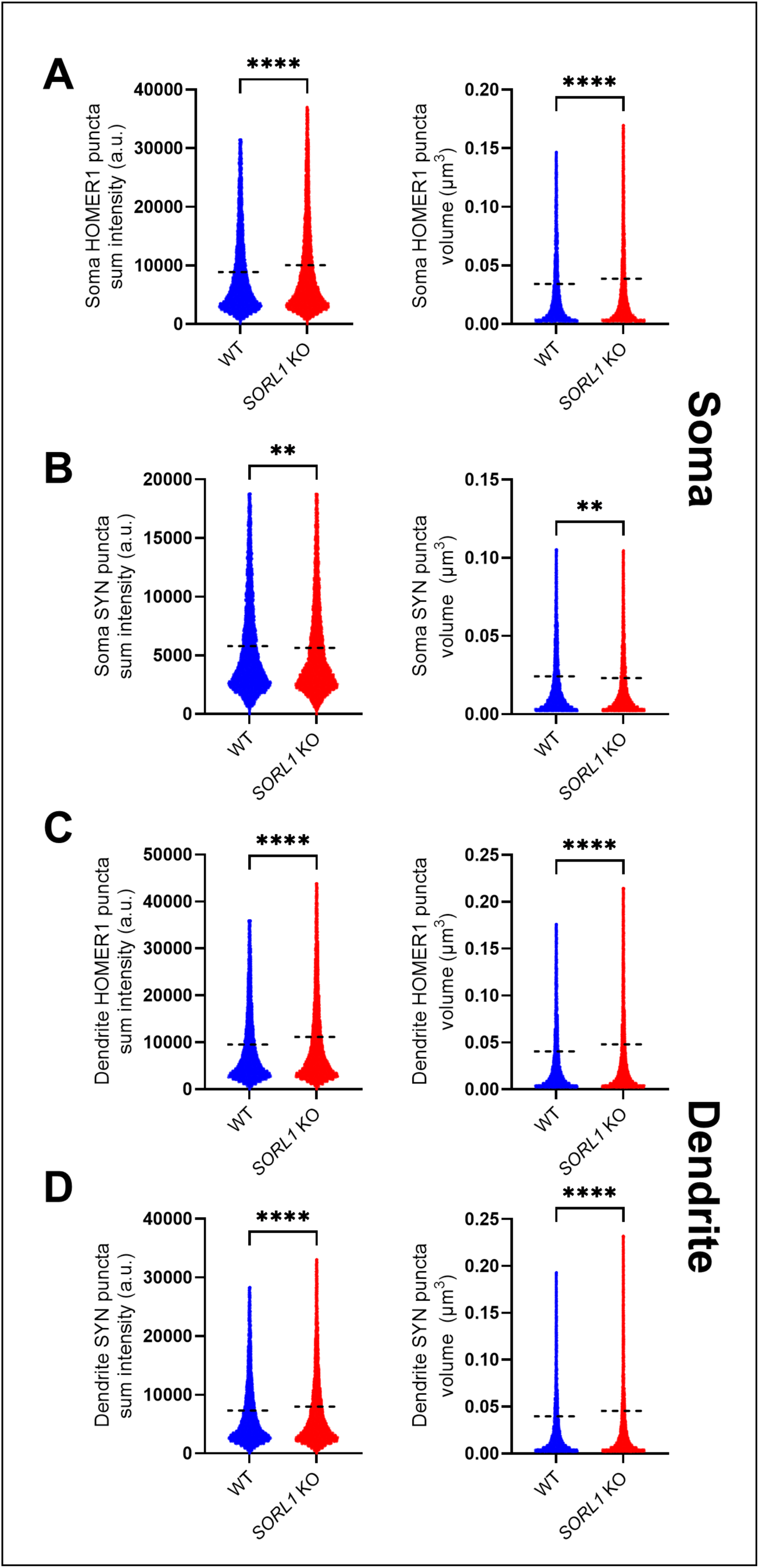
Abnormal *SORL1* KO postsynaptic densities are localized to the dendritic compartment. Quantification of pre and post synaptic SYNAPSIN and HOMER1 puncta in somatic (A,B) and dendritic (C,D) compartments of DPP 30 wild-type and *SORL1* KO neurons. A-B: Pre and post synaptic puncta volume and sum puncta fluorescence intensity in neuronal soma. C-D: Pre and post synaptic puncta volume and sum puncta fluorescence intensity in neuronal dendrites. HOMER1 puncta in the soma and dendrites of *SORL1* KO neurons were larger and contained greater fluorescence signal. Smaller SYNAPSIN puncta were detected in the soma of *SORL1* KO neurons with lower intensity but this difference did not extend into the dendrites, where greater volumes and intensities of presynaptic puncta in *SORL1* KO neurons were detected. 20,000 to 25,000 HOMER1 puncta and 12,000 SYNAPSIN puncta were identified and analyzed in somatic compartments. 60,000 to 100,000 HOMER1 puncta and 30,000 to 50,000 SYNAPSIN puncta were identified and analyzed in dendritic compartments. N = 32-34 images per genotype. Two clones per genotype were analyzed. Puncta data points plotted with mean values indicated by horizontal dashed line. Outliers were identified using the ROUT (Q = 1%) method and removed for statistical testing and plotting. Data were tested for normality and were determined to be not normally distributed (D’Agostino and Pearson test). Statistical testing for all quantifications was the Kolmogorov-Smirnov test, with significance marked as follows: P < 0.01 (**), P < 0.0001 (****). Not significant comparisons are unmarked.

**Figure S2:**
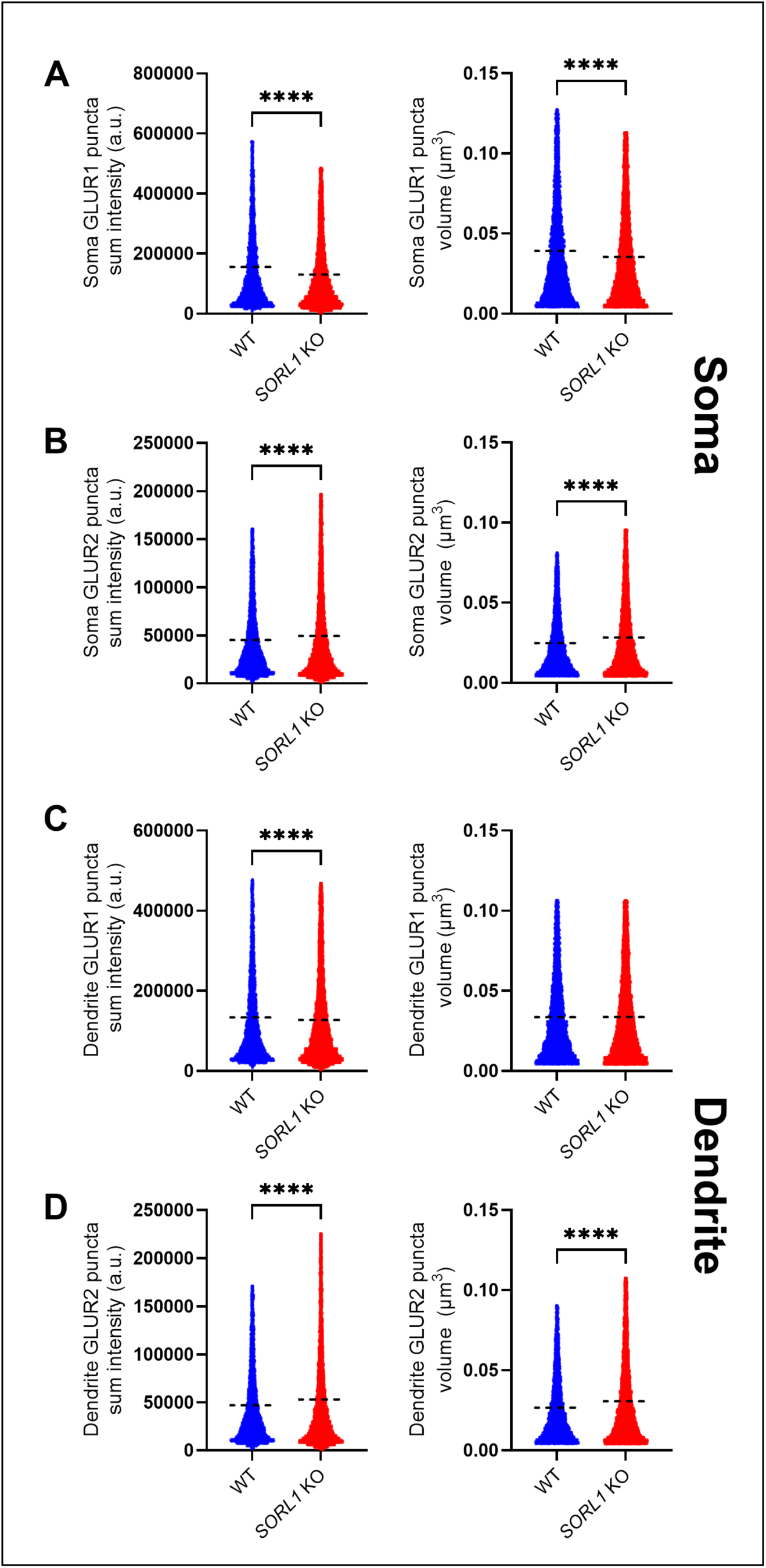
Wild-type and *SORL1* KO neuron GLUR1 and GLUR2 levels are consistent across soma and dendrites. Quantification of GLUR1 and GLUR2 immunofluorescent signal captured from matured DPP 30 wild-type and *SORL1* KO neuronal somatic and dendritic compartments. A-B: Soma localized GLUR1 and GLUR2 puncta sum intensity and volumes. C-D: Dendrite localized GLUR1 and GLUR2 puncta sum intensity and volumes in wild-type and *SORL1* KO neurons. With the exception of GLUR1 puncta volume in dendrites, differences of puncta sum intensity and volume match whole-cell analysis (Figure 3), indicating that changes in GLUR1 and GLUR2 are cell-wide and not localized imbalances of somatic and dendritic levels of the AMPA subunits. N = 10,000-12,000 GLUR1 and GLUR2 puncta were analyzed from somatic and dendritic neuronal structures. 32-34 images per genotype. Two clones per genotype were analyzed. Puncta data points plotted with mean values indicated by horizontal dashed line. Outliers were identified using the ROUT (Q = 1%) method and removed for statistical testing and plotting. Data were tested for normality and were determined to be not normally distributed (D’Agostino and Pearson test). Statistical testing for all quantifications was the Kolmogorov-Smirnov test, with significance marked as follows: P < 0.0001 (****). Not significant comparisons are unmarked

**Figure S3:**
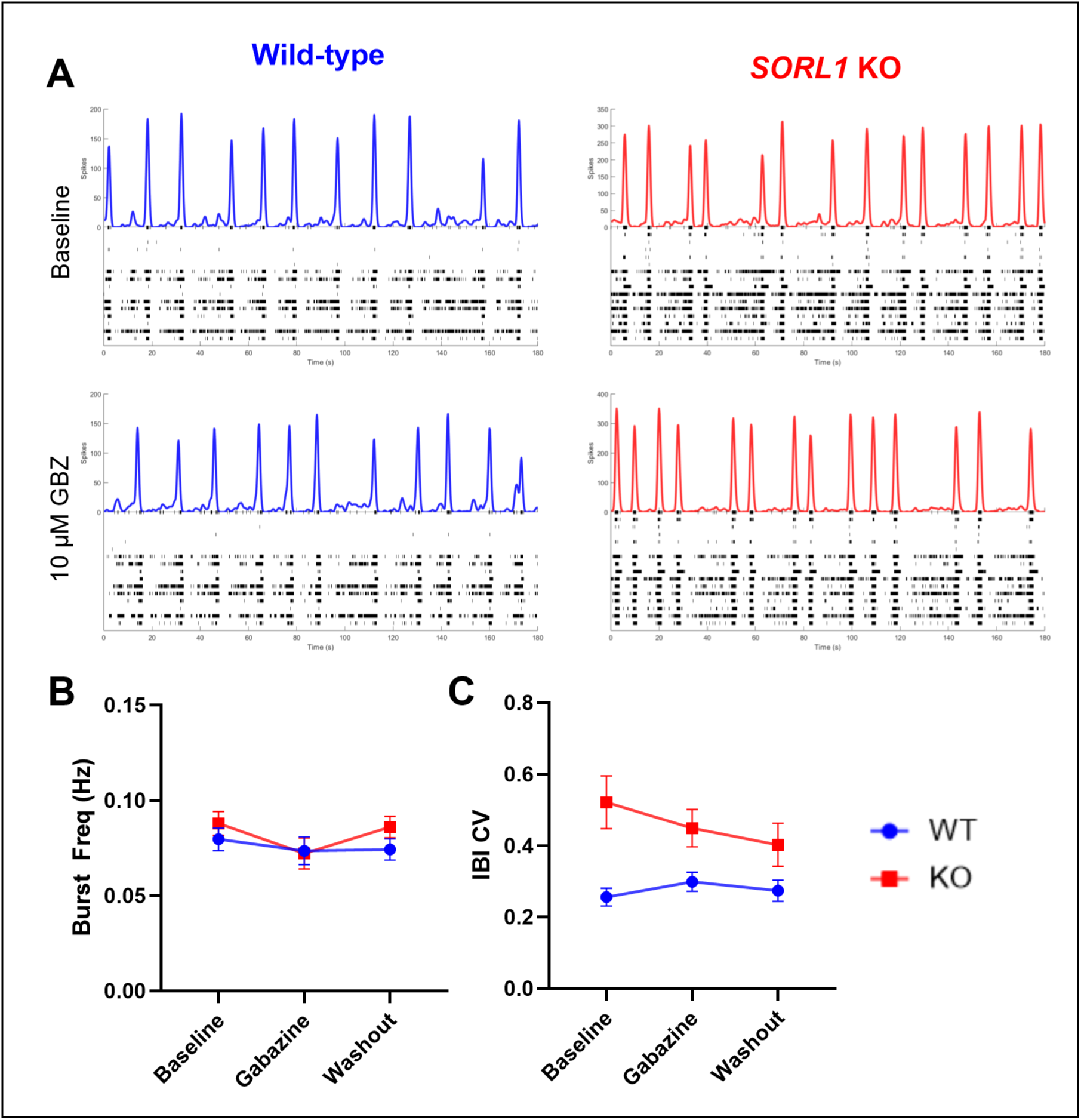
GABA_A_ mediated inhibitory neurotransmission does not contribute to *SORL1* KO hyperexcitability. A: Representative spike histograms (top panels) and raster plots (bottom panels) of wild-type and *SORL1* KO MEA cultures treated with GABA_A_ receptor antagonist Gabazine (10 μM). Little to no change is observed with Gabazine treatment, suggesting that inhibitory neurotransmission is not influencing or shaping patterns of electrical activity in these cultures. B: Network bursting rate of wild-type and *SORL1* KO cultures with GABA_A_ blockade. No changes are observed in wild-type or *SORL1* KO cultures. C: Coefficient of variation of network burst inter-burst intervals (IBI CV). Two-way ANOVA analysis revealed no significant interaction effect of genotype and Gabazine condition. N = 20-21 wild-type and *SORL1* KO cultures were used for Gabazine experiments at approximately DPP 45. Two wild-type clones and one *SORL1* KO clone were utilized. Data are represented by mean value ± SEM. Significance values of two-way ANOVA interaction results (cell line X drug condition) were not significant for both network bursting rate and IBI CV.

**Figure S4:**
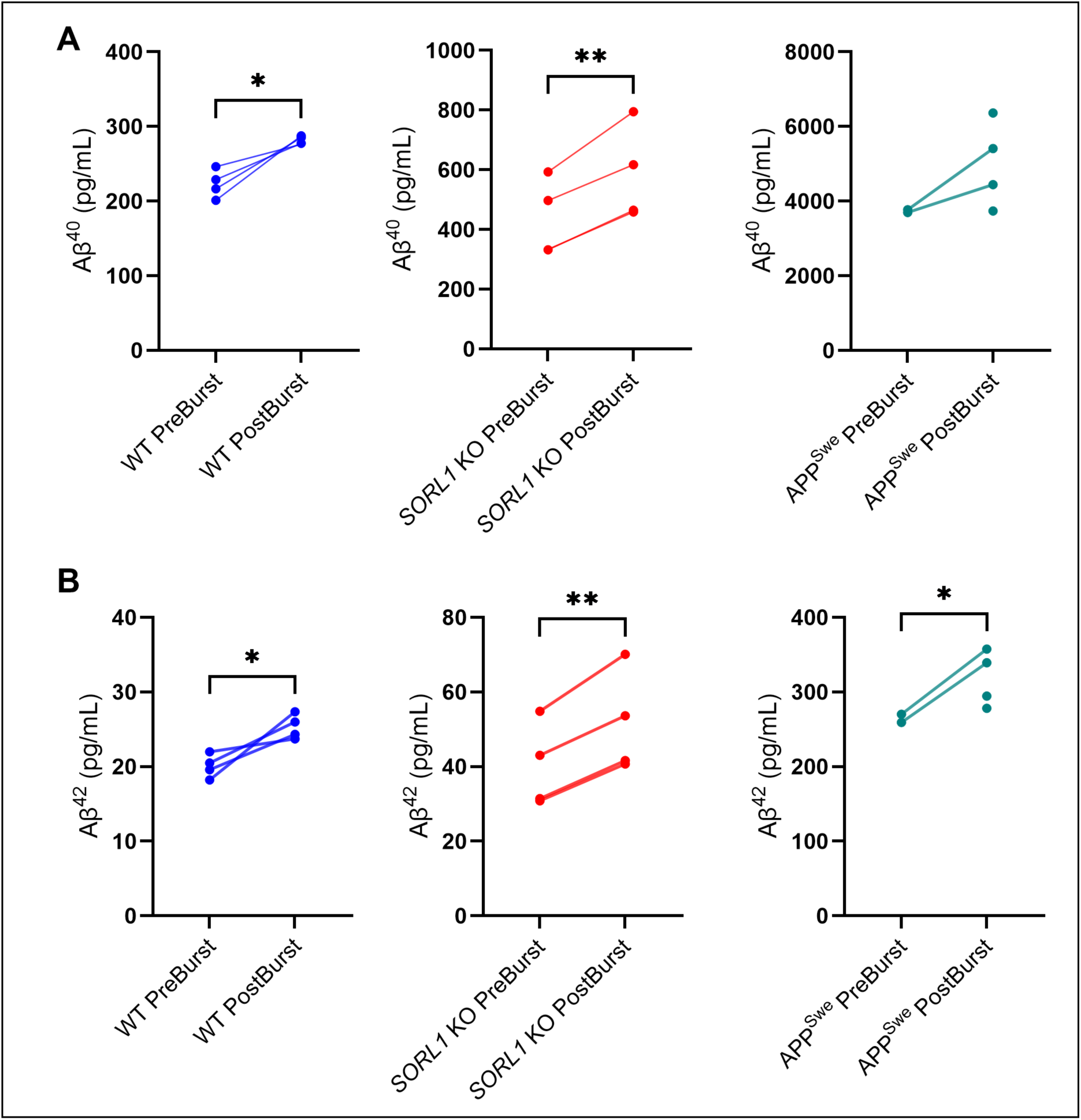
Amyloid beta secretion increases with neuronal network bursting. A: Concentrations (pg/ml) of secreted Aβ 1-40 harvested from wild-type, *SORL1* KO, and APP^Swe^ neurons before and after the onset of network bursting (DPP 14 and DPP 35, respectively). Wild-type and *SORL1* KO neurons exhibited a significant increase in Aβ^1-40^ secretion after the onset of synchronized network bursting. APP^Swe^ neurons change in Aβ^1-40^ trended upwards as well. B: Concentrations of (pg/ml) of secreted Aβ 1-42 harvested from wild-type, *SORL1* KO, and APP^Swe^ neurons before and after the establishment of synchronized network bursting (DPP 14 and DPP 35, respectively). All three cell lines secreted significantly more Aβ^1-42^ following network bursting onset. N = 4 wild-type and *SORL1* KO cultures. N = 2 APP^Swe +/+^ cultures. 2 clones of wild-type and *SORL1* KO cell lines were used, 1 clone of APP^Swe +/+^ was used. Each point represents the average of two technical replicates measured from conditioned medium collected from one MEA well ± SD. Paired t-tests results are indicated as follows: P ≤ 0.05 (*) and P ≤ 0.01 (**). Not significant comparisons are not marked.

**Table S1.**
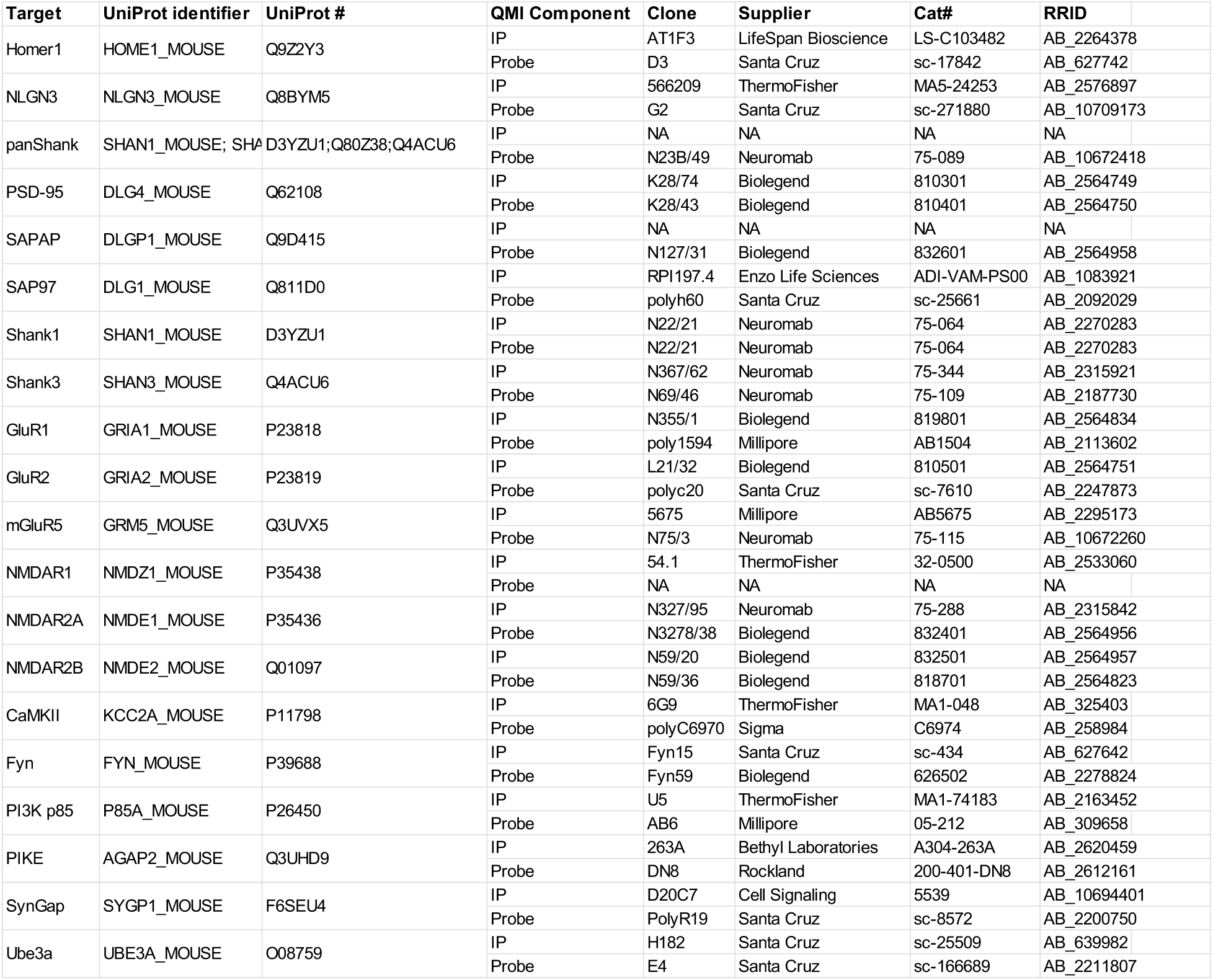

